# Widespread prevalence of a post-translational modification in activation of an essential bacterial DNA damage response

**DOI:** 10.1101/2023.10.09.561495

**Authors:** Aditya Kamat, Ngat T. Tran, Mohak Sharda, Neha Sontakke, Tung B. K. Le, Anjana Badrinarayanan

## Abstract

DNA methylation plays central roles in diverse cellular processes, ranging from error-correction during replication to regulation of bacterial defense mechanisms. Nevertheless, certain aberrant methylation modifications can have lethal consequences. The mechanisms by which bacteria detect and respond to such damage remain incompletely understood. Here, we discover a highly conserved but previously uncharacterized transcription factor (Cada2), which orchestrates a methylation-dependent adaptive response in *Caulobacter*. This response operates independently of the SOS response, governs the expression of genes crucial for direct repair, and is essential for surviving methylation-induced damage. Our molecular investigation of Cada2 reveals a cysteine methylation-dependent post-translational modification and mode of action distinct from its *E. coli* counterpart, a trait conserved across all bacteria harboring a Cada2-like homolog instead. Extending across the bacterial kingdom, our findings support the notion of divergence and co-evolution of adaptive response transcription factors and their corresponding sequence-specific DNA motifs. Despite this diversity, the ubiquitous prevalence of adaptive response regulators underscores the significance of a transcriptional switch, mediated by methylation post-translational modification, in driving a specific and essential bacterial DNA damage response.

## Introduction

Information regarding biological processes is primarily encoded in the genomes of living organisms. Functional modalities are further enhanced via modulation of the epigenetic states of DNA [1–4]. For example, methylation of DNA occurs in organisms across all domains of life. In bacteria, physiological DNA methylations at N5-meC, N4-meC and N6-meA nucleotide positions play diverse functions ranging from defensive roles involving immunity against invasive genetic elements to regulatory roles in cell cycle and transcriptional control [2,5]. In contrast to these physiological modifications, certain types of methylations can be aberrant. DNA methylations on O6-meG and O4-meT positions act as a potent source of mutagenesis, while N3-meA, N1-meA and N3-meC block DNA replication and transcription [6–9].

Bacteria cope with this threat of aberrant methylations via eliciting two DNA damage responses (DDR). The first response is the well-characterized and ubiquitously conserved SOS response that consists of two regulatory units, RecA and LexA [10]. The transcriptional repressor LexA occupies *lexA* boxes present in the promoter regions of SOS genes and inhibits their expression. Upon DNA damage, an activated RecA nucleoprotein filament triggers the autocleavage of LexA leading to de-repression of the SOS response [11–13]. Three distinct features are hallmarks of this response a). It is induced under all forms of DNA damage and is not specific to methylation damage alone [14], b). The response has leaky expression even in the absence of damage [15,16] and c). The response can result in mutagenesis due to expression of translesion synthesis polymerases [17]. Thus, SOS response induction is a trade-off between its essentiality and its cost.

A temporally delayed DDR subsequent to the SOS response, known as the adaptive response to methylation damage (or the Ada response) has also been reported in *E. coli* [18–20]. This response exhibits key distinguishing features compared to the SOS response: a). It is activated independent of the SOS response and is specific to methylation damage only [21], b). The response is adaptive i.e., exposure to a sub-lethal dose of methylation damage encodes for memory that allows cells to adapt and survive formerly lethal levels of DNA methylation damage [18,21], c). The response exhibits bi-stability [22]; a subpopulation of cells do not induce the response even under continuous exposure to methylation damage, and d). Repair via this response is non-mutagenic [18,23].

The master regulator of this DDR is the methyltransferase EcAda, comprising of an AdaA domain in its N-terminus (which associates with ‘A’ and ‘B’ DNA boxes located upstream of EcAda-regulated promoters) and a C-terminal methyltransferase domain [19]. EcAda expression is tightly regulated, with only 0-2 molecules of Ada present in cells in the absence of damage [22]. Post-translational modification (PTM) via methylation of a conserved cysteine in N-Ada is crucial for activating EcAda as a transcription factor [24–26]. Once activated, EcAda drives its own expression as well as expression of genes involved in direct repair of methylation lesions.

How prevalent is the risk of methylation damage, which could have resulted in the evolution of a methylation-specific DDR? Instances of encountering methylation DNA damage are not infrequent, and bacteria often face methylation stress. Intrinsic factors, such as insidious metabolic byproducts (e.g., lipid peroxidation) pose a prominent risk for DNA methylation damage [9,19]. Environmental factors including halocarbons and methylating agents produced by bacteria as instruments of inter-microbial warfare also contribute to this damage [27,28]. Additionally, infected mammalian cells subject invading bacteria to methylation stress [29]. More recently, studies have reported instances where physiological DNA methyltransferases, which are otherwise innocuous, erroneously inflict methylation damage on genomic DNA [30]. Thus, given the prevalence of methylation stress, it is likely that bacteria have evolved unique systems such as the Ada response to protect their genomes.

Intriguingly, despite the simultaneous discovery of the adaptive response to methylation damage alongside the SOS response [18,31], our knowledge of this pathway is presently restricted only to the *E. coli* paradigm. Thus, the significance and conservation of a methylation-based PTM in the regulation of a damage-specific bacterial DDR remains unknown. Indeed, limited number of computational studies suggest only sporadic conservation of EcAda across the bacterial kingdom [32–34], and many bacteria, such as *Caulobacter crescentus,* are reported to lack an inducible adaptive response altogether [35,36].

In this work, we discover a methylation-specific DDR in *Caulobacter crescentus*, which showcases key features of an adaptive response. This pathway is regulated by a conserved but uncharacterized transcription factor *ccna_03845* (‘Cada2’ henceforth). Detailed molecular characterization reveals a novel sequence-specific DNA-binding domain in this protein, as well as a methylation-based PTM required for activation of Cada2 as a transcription factor in a manner that is distinct from its *E. coli* counterpart. Despite the contrasting mechanistic features of EcAda and Cada2, we observe remarkable similarities in the defining characteristics of the downstream response. Phylogenetic distribution of adaptive response regulatory proteins further reveals their widespread prevalence across the bacterial kingdom, with ubiquitous conservation of key residues required for methylation-based activation (either in an EcAda-like or Cada2-like form). Collectively, our work highlights the importance of a transcriptional switch mediated by methylation post-translational modification in activating an essential bacterial response to methylation DNA damage.

## Results

### A methylation-specific DNA damage response in Caulobacter

We undertook a comprehensive transcriptomic approach to identify whether *Caulobacter* induces an SOS-independent transcriptional response specific to methylation damage. For this, we treated *Caulobacter* cells with agents that predominantly induce one specific form of DNA damage (methyl methane sulphonate (MMS) a causative agent of methylation DNA damage [37], mitomycin-C (MMC) that induces intra-strand crosslinks and mono-adducts [38], and norfloxacin that induces double-strand breaks [39]). We collected samples at 0, 20 and 40 minutes post damage exposure for RNA sequencing (Fig. 1A, Fig. S1A). Our analysis revealed several genes that were induced in all conditions (‘universal’) and a set of genes that were specifically induced under MMS treatment, but in none of the other damaging conditions (methylation-specific) (Fig. S1A).

**Figure 1.**
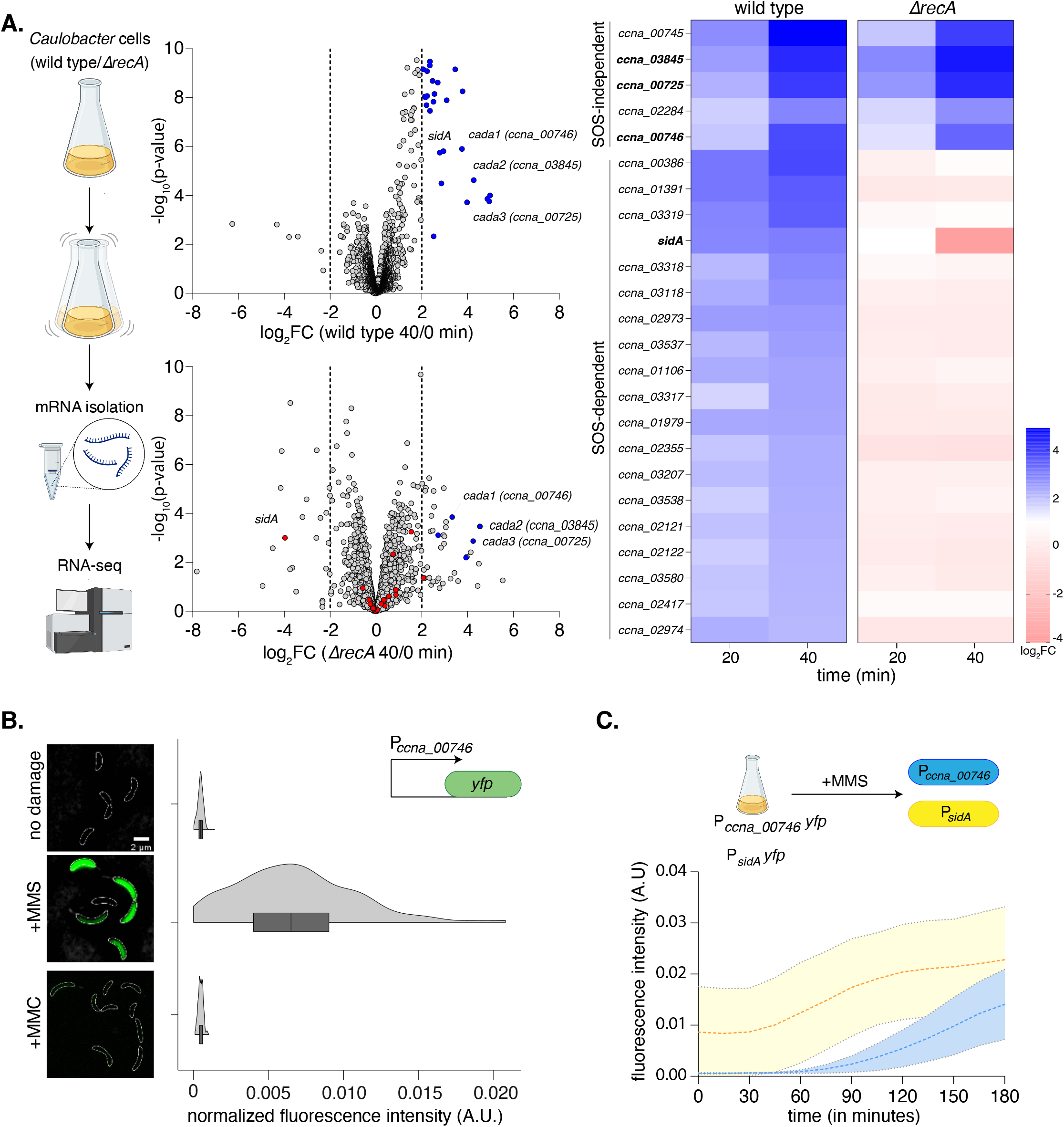
A methylation-specific DNA damage response in Caulobacter. **(A)** [Left] Schematic summarizing the RNA–sequencing experiment is provided. Wild type and *ΔrecA* (SOS-deficient) cells were exposed to 1.5 mM methyl methanesulphonate (MMS) for 40 minutes. [Centre] Volcano plots representing differentially expressed genes in comparison to cells exposed to no damage for wild type and *ΔrecA* cells respectively. Genes upregulated in wild type cells (log_2_FC > 2 and – log_10_(p-value) > 2) are highlighted in blue, while genes highlighted in red represent genes upregulated in wild type but downregulated in *ΔrecA* cells. [Right] Heat maps represent log_2_ FC values for individual genes induced in wild type and *ΔrecA* cells under MMS exposure. **(B)** Wild type P*_ccna_00746_-yfp* reporter (schematic inset) was exposed to 1.5mM MMS or 0.5μg/ml MMC respectively for 2 hours. Representative cells are shown on the left (scale bar: 5 μm, cell boundaries are marked by white dotted outline, here and in all other images). Violin plots show fluorescence intensity distribution normalized to cell area from single cells (n=300, from three biological replicates) **(C)** [Top] Schematic of the experimental protocol comparing induction kinetics of the *Caulobacter* SOS and methylation-specific response via time lapse microscopy. [Bottom] Fluorescence intensity normalised to cell area for P*_ccna_00746_-yfp* and P*_sidA_-yfp* cells over 3 hours of exposure to 1.5mM MMS.

We next asked whether the expression of these genes was independent of the SOS response. We subjected *recA* deleted cells to MMS treatment and carried out RNA sequencing in this background. Comparing wild type to *recA* deletion cells showed that the methylation damage-specific genes were induced in a *recA*-independent manner (SOS-independent) (Fig. 1A). In contrast, genes that were induced under all DNA damaging conditions in wild type background remain uninduced in *recA* deletion cells (SOS-dependent) (Fig. 1A, Fig. S1A).

We identified *ccna_00745* to be the highest induced under methylation damage in the transcriptome analysis (Fig. 1A). This gene is co-operonic with a second gene *ccna_00746,* that is also induced in a methylation DNA damage-specific manner (Fig. S1A). To further corroborate the RNA-seq observations, we constructed a fluorescence-based readout for promoter activity of these genes (P*_ccna_00746_-yfp*). No significant expression of *yfp* was detected in the absence of damage (Fig. 1B). We observed YFP expression from this construct only in cells treated with MMS, and not other DNA damaging agents MMC, Norfloxacin and hydroxyurea (HU) (Fig. 1B, Fig. S1B). We additionally utilized a second methylation damaging agent, streptozotocin (STZ), a naturally occurring antibiotic produced by *Streptomyces achromogenes* var. *streptozoticus* [27,28]. In this case as well, we observed YFP expression from the methylation damage-specific promoter (Fig. S1B). These observations were distinct from those seen for a fluorescence reporter for the promoter of *sidA,* a known SOS-dependent gene [40], that has been reported to be induced under other types of DNA damage in a RecA-dependent manner [40,41]. Thus, *Caulobacter* elicits a transcriptional response to methylation damage that is independent of the SOS response.

We next investigated the temporal kinetics of the methylation-specific response in relation to the SOS response. For this, we carried out time-lapse imaging of cells carrying either the P*_ccna_00746_-yfp* or the P*_sidA_-yfp* reporter after MMS exposure. We observed that expression of *yfp* from the methylation damage-specific promoter expression was delayed when compared to the SOS reporter, and succeeded the SOS response (Fig. 1C). Finally, we tested whether the induction kinetics of this response is adaptive. For this, we first exposed our promoter fusion strain to a low (0.5mM) dose of MMS. This led to only a modest increase in the expression of P*_ccna_00746_-yfp* (Fig. S1C). Consistent with an ‘adaptive’ response [42] these pre-treated cells were able to induce P*_ccna_00746_-yfp* significantly faster as compared to untreated cells upon exposure to a higher dose (1.5mM) of MMS (Fig. S1C). Together, the *Caulobacter* methylation damage-specific response showcases key features of an adaptive response to methylation damage as reported in *E. coli*. Based on these observations, we henceforth refer to this response as the *Caulobacter* adaptive response to methylation damage.

### Caulobacter adaptive response to methylation damage is regulated by Cada2

Given the striking similarity between the adaptive responses of *Caulobacter* and *E. coli,* we wondered whether the *Caulobacter* response was regulated by an EcAda-like protein [43]. While we could not identify any EcAda-like protein in *Caulobacter*, we observed that three adaptive response candidates (*ccna_00746*, *ccna_03845* and *ccna_00725*) possessed an Ada-like methyltransferase domain (PF01035) in their C-terminus region (Fig. 2A). Intriguingly, domains corresponding to N-Ada of EcAda protein (required for forming sequence-specific interactions with the cognate Ada promoters) were split between *ccna_00746* (with A box-binding domain) and *ccna_03845* (with B box-binding domain) (Fig. 2A). We thus annotated these as ‘*cada’* (***C****aulobacter **ada**ptive response*) genes *cada1*, *cada2* and *cada3*.

**Figure 2.**
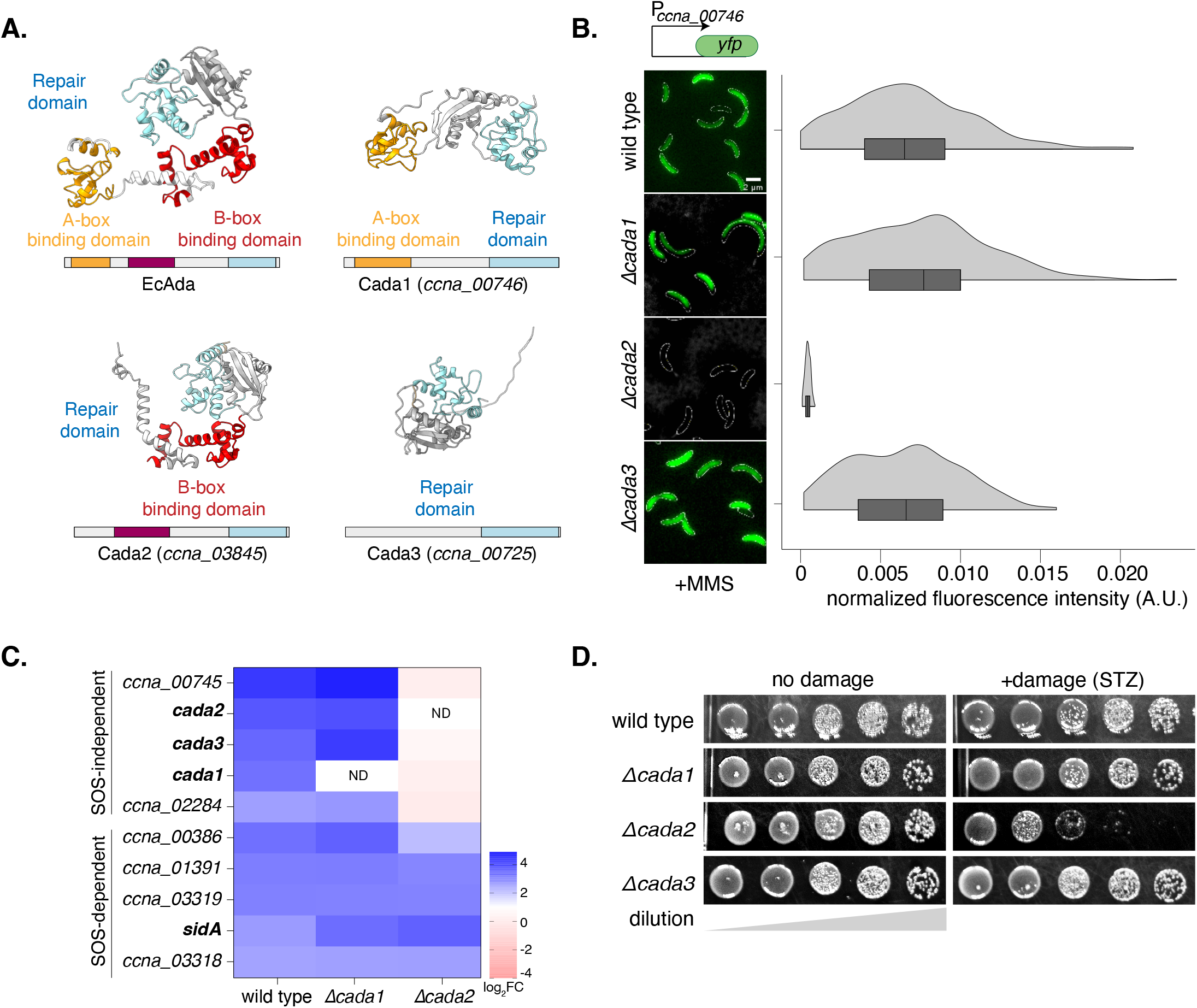
Caulobacter adaptive response to methylation damage is regulated by Cada2. **(A)** Comparative analysis of high-confidence Alphafold structural models of EcAda, Cada1, Cada2 and Cada3 proteins along with their domain organizations. **(B)** [Left] Representative cells showing P*_ccna_00746_-yfp* reporter induction in wild type (from Fig. 1B) and individual deletion strains of the *cada* genes under 1.5mM MMS damage. [Right] Violin plots showing fluorescence intensity distribution normalized to cell area from single cells (n=300, from three biological replicates). **(C)** Heat map of log_2_FC values for genes upregulated in wild type, *Δcada1* and *Δcada2* cells following 1.5 mM MMS treatment. **(D)** Survival assay of individual deletions of *cada* genes with and without methylating agent (STZ) (5μg/ml).

Next, we asked whether Cada1, Cada2 or Cada3 regulated the adaptive response. We deleted all three *cada* genes individually and assessed the activity of the P*_ccna_00746_-yfp* reporter. We found that only cells lacking *cada2* were unable to induce P*_ccna_00746_-yfp* under methylation damage (Fig. 2B). In support, RNA-sequencing analysis revealed that other MMS-specific, RecA-independent genes were also not induced in cells lacking *cada2*, but were unaffected in cells lacking *cada1* (Fig. 2C). Thus, A-box containing Cada1 does not drive gene expression under the adaptive response, while Cada2 possessing solely the B-box binding domain is required for response activation. Cell survival upon exposure to the natural antibiotic streptozotocin was also significantly compromised in cells lacking *cada2* (Fig. 2D), suggesting that the Cada2-dependent adaptive response to methylation damage was an essential DDR pathway in *Caulobacter*.

### Cada2 associates with adaptive response promoters in a sequence-specific manner

To determine whether Cada2 was directly responsible for the induction of the adaptive response genes under methylation damage, we performed ChIP-sequencing experiments (Fig. 3A). For this, cells expressing *cada2-3x-flag* from its endogenous promoter were used. The flag-tagged strain resembled survival of wild type cells under methylation DNA damage and damage-dependent expression of Cada2 was detected via western blot (Fig. S2A-B). Using this strain, we carried out ChIP-seq experiments in the presence or absence of methylation damage (Fig. 3A). We estimated Cada2-3xFlag enrichment across the *Caulobacter* genome under damage using untagged wild type cells as control. This allowed us to identify bona fide Cada2-binding sites.

**Figure 3.**
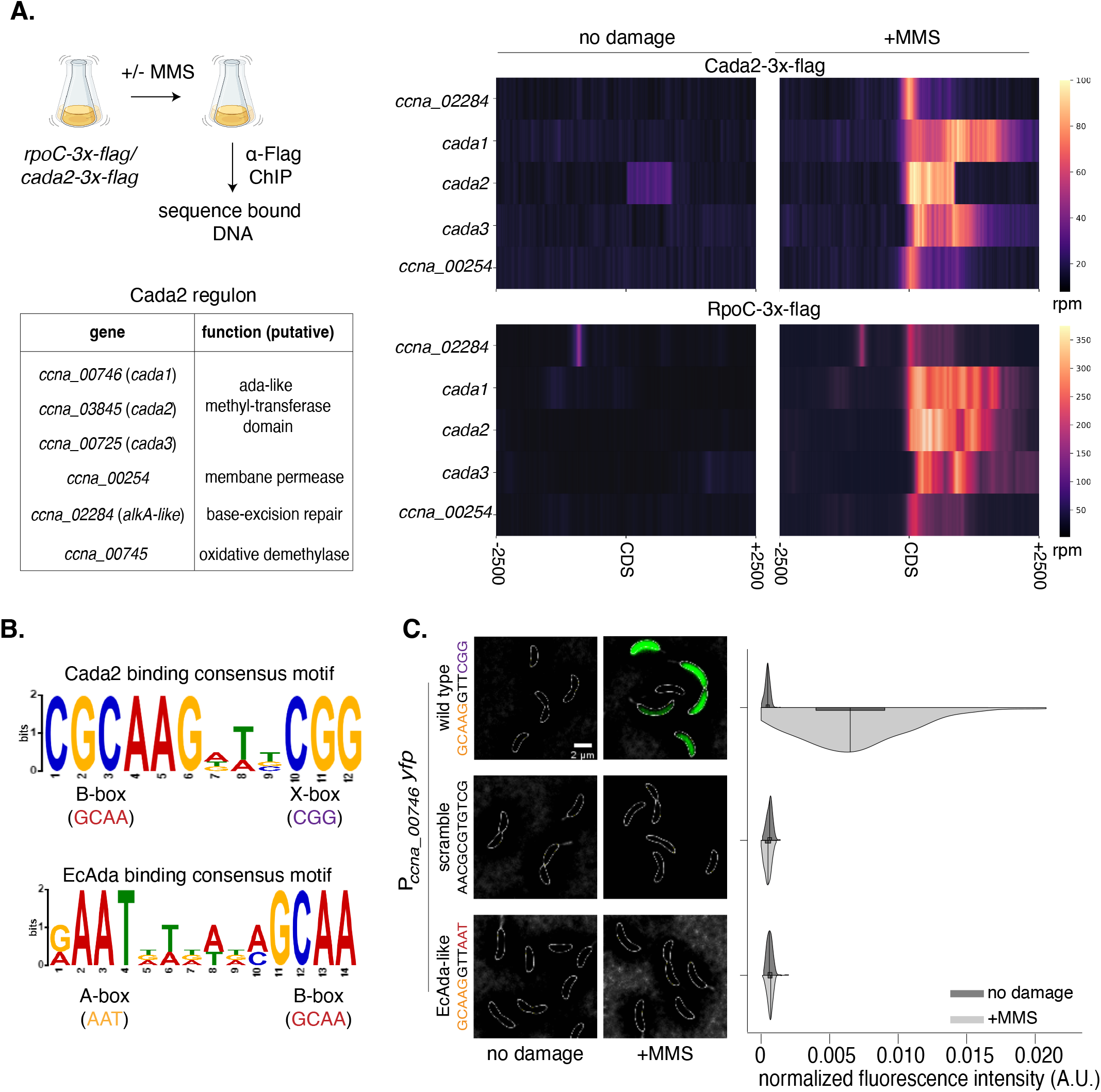
Cada2 associates with adaptive response promoters in a sequence-specific manner. **(A)** [Top left] Schematic of the ChIP-sequencing protocol. [Bottom left] Table showcasing the Cada2 regulon. Gene names and predicted function are listed. [Top right] Normalised reads (in rpm) for *Cada2-3X-Flag* ChIP-seq represented ±2.5kb around the CDS of the Cada2 regulon genes in the presence/absence of 1.5mM MMS damage. [Bottom right] Data represented in a similar fashion for *RpoC-3X-Flag* ChIP-seq. **(B)** Binding consensus motif for Cada2 and EcAda derived from the promoters of the respective constituents of their regulon. **(C)** Representative cells showing induction of variants of the P*_ccna_00746_-yfp* reporter (scramble/*EcAda*-like) compared to wild type (from Fig. 1B) under 1.5mM MMS damage. [Bottom right] Half-violin plots show fluorescence intensity distribution normalized to cell area from single cells for the reporter variants in the presence (dark) and absence of damage (light) (n=300, from three biological replicates).

Under damage, we observed Cada2 enrichment at its own promoter as well as at other promoters of the adaptive response (Fig. 3A, Fig. S2C-D). This autoregulation of its own expression appears to be a conserved feature between EcAda [44] and Cada2. To further assess the methylation damage-dependent transcription of these genes, we performed ChIP-seq with flag-tagged RNA polymerase subunit C (RpoC-3xflag) construct. We did not observe localization of RpoC to the *cada2-*associated promoters in the absence of methylation damage (Fig. 3A, Fig. S2C-D). Upon exposure to damage, RpoC signal could be detected at the *cada2* promoter, as well as other adaptive response promoters (Fig. 3A, Fig. S2C-D). ChIP-seq results overlapped significantly with the Cada2-dependent upregulated genes identified via RNA-seq manner (∼83%). Superimposing the two datasets allowed us to describe the Cada2 regulon comprising six genes (Fig. 3A), most of which appear to have direct repair-associated activities.

We carried out MEME analysis of Cada2-bound regions to identify any sequence-specific DNA binding motifs. This revealed two sequences consistent across all promoters of the *Caulobacter* adaptive response (Fig. 3B). One of these sequences, ‘GCAA’, was identical to the B-box sequence motif bound by the B-box binding domain of EcAda (Fig. 3B). Thus, we annotate it as the B-box motif in case of *Caulobacter* as well. We did not detect an A-box sequence motif (‘AAT’, bound by EcAda A-box binding domain). We instead identified a second recurrent motif that was GC-rich (‘CGG’). We annotated this as ‘X-box’ sequence motif. These two sequence motifs were separated by a 3-base pair spacer which exhibited no recurrently conserved sequence, but seemed to be consistently AT-rich. Together, we annotated the B-box and X-box motif, separated by the AT-rich spacer as the Cada2 binding sequence motif (Fig. 3B).

To assess whether this sequence is required for Cada2-mediated regulation, we scrambled the complete sequence in our P*_ccna_00746_-yfp* reporter construct. We found that such a reporter was no longer induced under methylation damage (Fig. 3C). We next asked whether an EcAda-like DNA binding motif comprising of an ‘A-box’ instead of the ‘X-box’ DNA motif could drive promoter activity by Cada2. Here too we observed that our reporter construct carrying an A and B box DNA motif was not responsive to methylation damage (Fig. 3C). This suggests that the newly identified B+X-box motif is essential for the induction of the *Caulobacter* adaptive response.

### Cada2-like proteins are widespread and encode a novel DNA binding domain

The absence of the A-box DNA motif as well as the A-box protein domain in case of Cada2 led us to hypothesize that the X-box DNA motif must likely have a cognate DNA binding region in the Cada2 protein. We thus performed a comprehensive computational search for all Cada2-like methyltransferases across the bacterial kingdom. For this, we built a Hidden Markov Model (HMM) [45] profile from a multiple-sequence alignment of Cada2-like proteins. As comparison, we followed the same process to identify EcAda-like [19,26] and AdaA-like [46] proteins as well. Phylogenomic distribution of these proteins across a curated, non-redundant database of diverse bacterial species showcased that they are abundant and widespread across all major bacterial phyla (Fig. 4A and Fig. S3D). Furthermore, Cada2-like proteins infrequently co-occurred with EcAda-like proteins in the same genome (∼7%).

**Figure 4.**
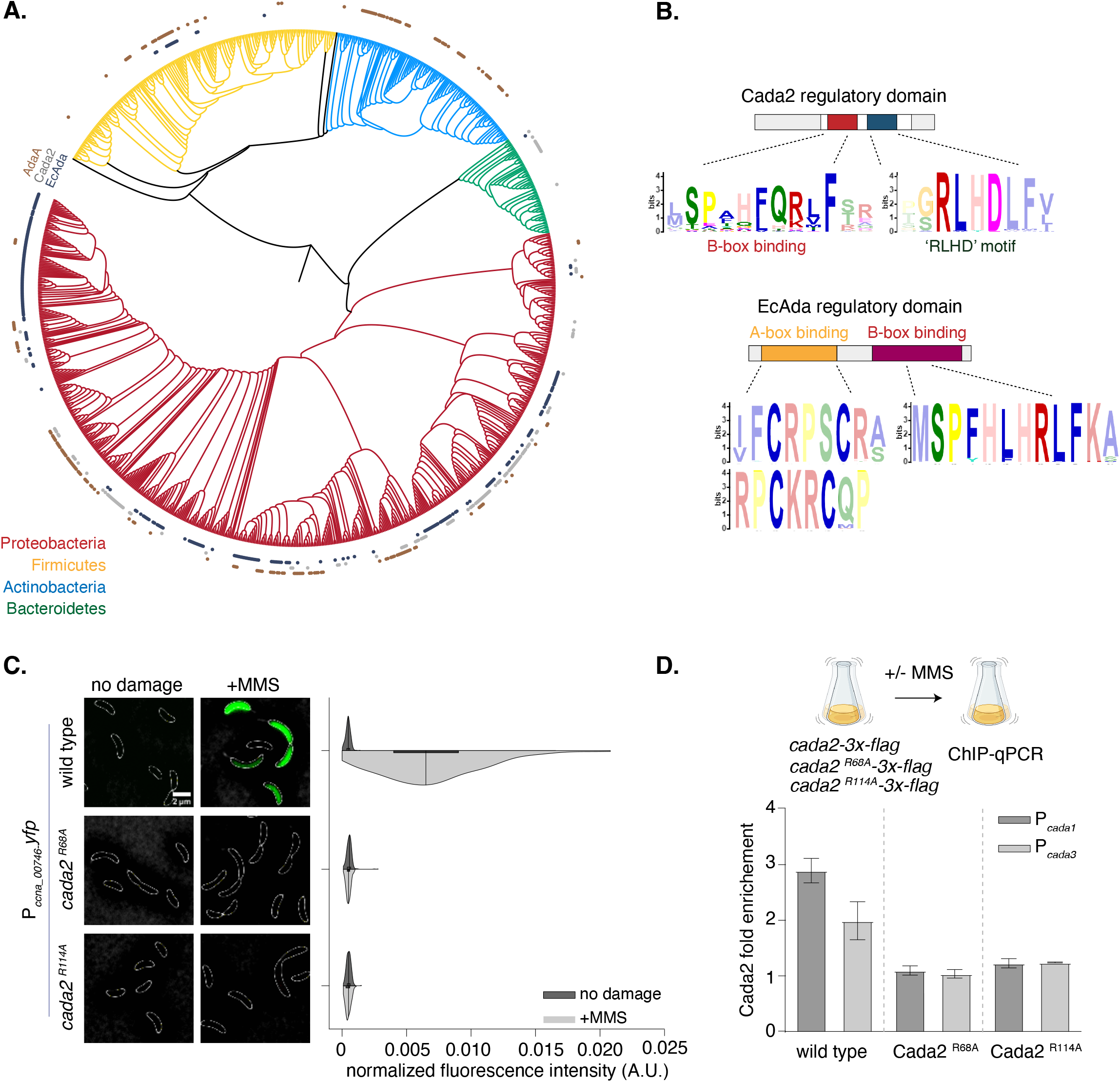
Cada2-like proteins are widespread and encode a novel DNA binding domain. **(A)** Phylogenetic distribution at the genus level of EcAda-like (black), Cada2-like (grey) and AdaA-like (brown) proteins across the genomes of four bacterial phyla: proteobacteria (red), firmicutes (yellow), actinomycetes (blue), bacteriodetes (green). Presence/ absence is shown on a 16S rRNA-based phylogenetic tree of bacterial genomes **(B)** [Top] MEME motif derived from the regulatory domain of 1000 EcAda-like proteins. [Bottom] MEME motif derived from 1000 Cada2-like proteins reveal conserved residues in the regulatory domain. **(C)** [Left] Representative images showing P*_ccna_00746_-yfp* reporter induction in wild type (from Fig. 1B) and *cada2* mutant (c*ada2^R68A^and* c*ada2^R114A^*) background under 1.5mM MMS damage. [Right] Half-violin plots show fluorescence intensity distribution normalized to cell area from single cells in the presence (dark) and absence of damage (light) (n=300, from three biological replicates). **(D)** ChIP-qPCR comparing wild type Cada2, Cada2^R68A^ and Cada2^R114A^ enrichment at *cada1* and *cada3* promoters following exposure to 1.5mM MMS. In all cases, flag-tagged version of protein is expressed from a xylose-inducible promoter on a high-copy replicating vector (in a *Δcada2* background). Bar graphs represent mean fold enrichment in comparison to wild type *cada2* (n=3 independent repeats, error bars represent standard error).

We zoomed into the DNA binding (regulatory) regions of EcAda and Cada2-like proteins to identify conserved and unique features. As anticipated, both classes of proteins encoded the B-box binding domain (marked by the conserved ‘SPHFQR’ amino acid residues (Fig. 4B)). It is likely that this region associates with the B-box DNA motif, that can be found in both EcAda and Cada2 regulons. The two proteins deviated with regards to the A-box, with A-box (marked by the four cysteine residues) being conserved in all EcAda-like proteins (Fig. 4B), but absent in all Cada2-like proteins. Instead, proximal to the putative B-box binding domain, Cada2-like proteins possessed a unique and highly conserved ‘RLHD’ sequence domain (Fig. 4B). We highlight here that the conservation of the RLHD domain is even more than that of the B-box. AlphaFold [47,48] model of the Cada2 protein predicted that this RLHD domain falls in a helix-turn-helix domain similar to the B-box binding domain (Fig. S3A).

In support of the significance of these putative DNA binding regions of Cada2, mutation of the conserved arginine (usually associated with DNA-binding) in both domains to an alanine residue (B-box *cada2^R68A^* and RLHD-motif *cada2^R114A^*) abrogated P*_ccna_00746_-yfp* reporter activity under methylation damage (Fig. 4C). The two mutants were not compromised in expression and phenocopied the *cada2* deletion when assessed for survival under methylation damage (Fig. S3B-C). Promoter binding was also compromised in both mutants as seen from ChIP-qPCR experiments for Cada2 association at promoter regions of genes belonging to the Cada2 regulon (Fig. 4D). We conclude that the B-box and the novel and conserved ‘RLHD’ domains on the Cada2 protein enable it to associate with its promoters in a sequence-specific manner.

### Cada2 is a methylation-dependent transcription factor

We noticed in the ChIP profiles that Cada2 appeared to be modestly enriched at its own promoter even in no damage conditions (Fig. 3A, Fig. S2C-D). This was in contrast to RNA polymerase, that was observed to bind to the *cada2* promoter region only in the present of methylation damage (Fig. 3A, Fig. S2C-D). Indeed, *Caulobacter* induced the adaptive response only upon exposure to DNA methylation damage (Fig. 1B), and overexpression of *cada2* from a xylose-inducible promoter was insufficient to induce expression from the P*_ccna_00746_-yfp* reporter in the absence of damage (Fig. 5A).

**Figure 5.**
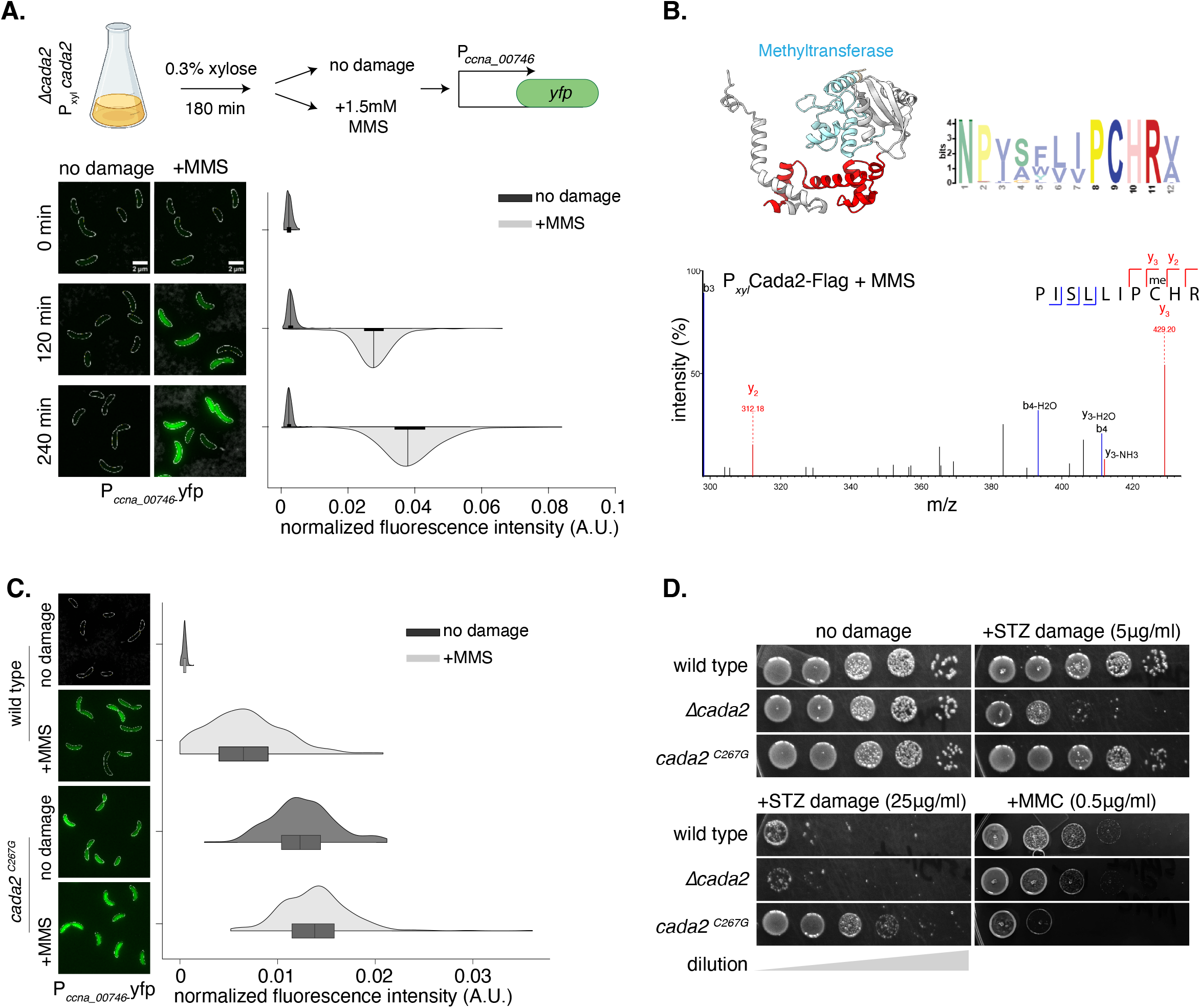
Cada2 is a methylation-dependent transcription factor. **(A)** [Top] Schematic representing experimental protocol for *cada2* overexpression analysis (see main text for details). [Bottom] Representative images and normalized fluorescence intensity for P*_ccna_00746_-yfp* expression where *cada2* is over-expressed from a xylose-inducible promoter in a *Δcada2* background in presence/ absence of 1.5mM MMS damage (n=300, from three biological replicates). **(B)** (Top left) Alphafold structure of Cada2 highlighting the presence of the methyltransferase domain is schematized. (Top right) MEME generated from 1000 bacterial Cada2 proteins indicate conservation of the ‘PCHR’ motif across methyltransferase domains. (Bottom) Zoomed in mass spectrometry fragmentation spectrum of a methylated peptide detected from a *cada2*-flag strain under 1.5mM MMS exposure. The peptide (inset) includes the conserved ‘PCHR’ motif of the Cada2 methyltransferase domain. Representative spectrum from 8 fragments across two biological replicates is shown. Methylation modification on Cys267 was detected in 6 out of 8 fragments. **(C)** [Left] Representative cells showing P*_ccna_00746_-yfp* reporter induction in wild type (from Fig. 1B) and c*ada2^C267G^* mutant under 1.5mM MMS damage. [Right] Violin plots show fluorescence intensity distribution normalized to cell area from single cells (n=300, from three biological replicates). (D) Survival assay of c*ada2^C267G^*mutant in the presence and absence of streptozotocin (5 and 25 μg/ml) and mitomycin C (0.5 μg/ml) damage.

We asked whether RNA polymerase recruitment to the promoter regions by Cada2 was mediated by physical interaction between Cada2 and the RNA polymerase holoenzyme, and if this step could be methylation-dependent. We thus carried out bacterial-two-hybrid interaction analysis of Cada2 against various RNA polymerase holoenzyme subunits (RpoA, RpoB, RpoC, RpoD and RpoZ). As a positive control we used the helicase-nuclease protein complex components AddA and AddB [49]. In the presence of methylation damage, we observed interaction signal between Cada2 and RNA polymerase subunit A (Fig. S4A).

How does methylation damage activate Cada2 function? Under damage, EcAda is post-translationally methylated at conserved cysteine residues in its A-box binding and methyltransferase (‘PCHR’) domains respectively [50,51]. However, it is the methylation of Cys_38_ in the A-box binding domain that is required for its activation as a transcription factor via modulation of sequence-specific DNA binding affinity of EcAda [26]. While Cada2 lacks the A-box binding domain, it does possess the methyltransferase domain (in its C-terminus) (Fig. 5B). Hence, we tested whether Cada2 is methylated post exposure to methylation DNA damage and whether this PTM is required for its activity.

Using mass spectrometry, we identified a methylation modification on a cysteine residue (Cys_267_, part of the ‘P**C**HR’ methyltransferase domain) of Cada2 in cells treated with methylation damage, with no detectable methylation in the absence of damage (Fig. 5B). We mutated the Cys_267_ residue to an alanine to disrupt the methylation modification. This mutant, *cada2^C267A^,* was unable to drive the induction of the P*_ccna_00746_-yfp* reporter and phenocopied a *cada2* deletion under streptozotocin treatment (Fig. S4B, Fig. S4C). Significantly, mutations in the promoter-binding domains of Cada2 (B-box *cada2^R68A^* and RLHD-motif *cada2^R114A^*) did not affect the ability of Cada2 to act as a methyltransferase, suggesting that this activity occurred in a DNA sequence-independent manner (Fig. S4D), and that methylation likely precedes the transcriptional response regulated by methylated Cada2.

To test if methylation of Cada2 was sufficient for its activation, we mutated the Cys_267_ residue to a glycine (*cada2^C267G^*). Such a mutation in the A-box of EcAda results in its constitutive activation as a transcription factor [26]. In contrast to the alanine mutant which abrogated *cada2* activity, we found that *cada2^C267G^* led to constitutive expression of *yfp* from the P*_ccna_00746_-yfp* reporter even in the absence of methylation damage (Fig. 5C). Furthermore, *cada2^C267G^* did not display growth defects associated with the *cada2* deletion or *cada2^C267A^* under methylation damage (Fig. S4C, Fig. 5D). Thus, Cada2 is activated as a regulator of the *Caulobacter* adaptive response via methylation of the conserved cysteine in its methyltransferase domain (PF01035).

Intriguingly, in comparison to wild type cells, the *cada2^C267G^*mutant had a significant growth advantage under methylation damage (Fig. 5D). However, sequence analysis of the ‘PCHR’ domain in Cada2-like proteins showed that a constitutive ‘ON’ version of this protein is not found in nature (Fig. 5B, 1000 non-redundant genomes analyzed). We thus wondered whether there could be growth conditions where such a response must remain repressed. To test this, we subjected wild type and *cada2^C267G^*mutant cells to MMC-treatment, an unrelated DNA damage condition where Cada2 activity is not essential and would ordinarily be in an ‘OFF’ state. In contrast to the growth advantage under methylation damage, we observed that *cada2^C267G^*cells were considerably compromised in growth in comparison to wild type when exposed to MMC (that induces mono-adducts and intra-strand crosslinks) (Fig. 5D). While the molecular mechanism underlying this crosstalk warrants further investigation, these observations support the possibility that a PTM-based methylation specific transcriptional switch is important for Cada2 regulation, due to antagonistic effects associated with this response being mis-regulated in other stress conditions.

### Regulatory features of Cada2 are conserved across bacteria and distinct from the EcAda paradigm

Taken together, the following regulatory features constitute the *Caulobacter* adaptive response: a. Promoter regions of genes under this regulon carry a B-box motif (GCAA) and an X-box motif (CGG). b. Cada2 protein N-terminus encodes B-box binding and RLHD domains, that enable it to interact with promoter regions in a sequence-specific manner. c. Cada2 protein C-terminus encodes a methyltransferase domain. Methylation of the conserved cysteine in this domain activates Cada2 as a transcription factor, likely via enabling interaction with RNA polymerase.

We asked how extrapolatable these observations would be across bacterial species. For this, we collated the presence or absence of the features listed above for a set of bacteria (including closely-related and unrelated) that carry either an EcAda-like or Cada2-like protein (Fig. 6A). To our surprise, we found that every organism that encoded an EcAda-like protein carried all the associated regulatory features for an *E. coli*-like adaptive response (Fig. 6A), while organisms with Cada2-like protein had regulatory features observed in case of the *Caulobacter* adaptive response (Fig. 6A).

**Figure 6.**
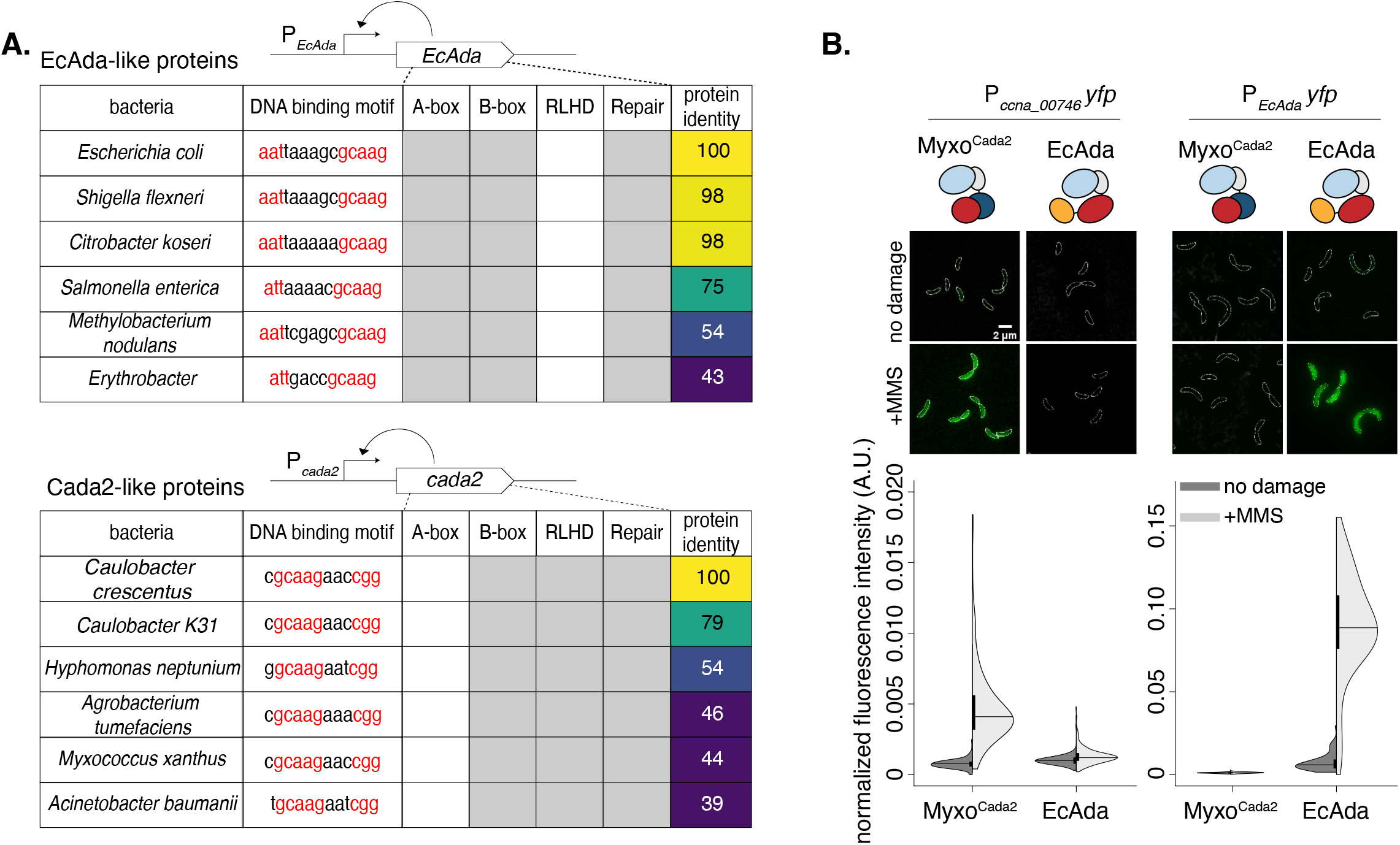
Regulatory features of Cada2 are conserved across bacteria and distinct from the EcAda paradigm. **(A)** Regulatory Ada methyltransferases such as EcAda and Cada2 are auto-regulatory; as represented in schematics, these proteins regulate activity of their own promoters (in a methylation-dependent manner). This allowed us to predict presence/absence of regulatory features in their respective promoters. The table represents conservation of regulatory features in the protein (A, B or RLHD amino acid sequences) and in the cognate promoter sequences (A, B or X Box DNA motif) of identified EcAda-like and Cada2-like proteins. Presence (grey) or absence of features (blank) is indicated along with % protein sequence. **(B)** Cada2 from *Myxococcus* (Myxo^Cada2^) can drive P*_ccna_00746_-yfp* induction in *Δcada2* in response to MMS damage, but cannot drive *yfp* expression from the *EcAda* promoter. Conversely, Ada from *E. coli* can activate expression of *yfp* from the *EcAda* promoter, but not from the P*_ccna_00746_* promoter.

The conservation of the protein and promoter sequence across Cada2-like proteins motivated us to test whether these proteins are functionally inter-changeable. For this, we expressed a Cada2-like protein from *Myxococcus xanthus* (Cada2^myxo^) in *Caulobacter* lacking its own *cada2*. Cada2^myxo^ has overall low identity (44%) to the *Caulobacter* Cada2 protein, however, it shares high similarity in terms of conserved regulatory features (Fig. 6A). As a control, we expressed EcAda in the same background to assess whether EcAda could complement the absence of Cada2. We found that Cada2^myxo^ was able to fully complement the absence of *Caulobacter cada2* under methylation damage, as assessed by cell survival as well as promoter activity of P*_ccna_00746_-yfp* (Fig. 6B, Fig. S5A). In contrast, EcAda was not able to complement a *cada2* deletion phenotype (Fig. 6B). This protein, expressed in *Caulobacter,* was fully proficient in driving expression of *yfp* from a promoter carrying the EcAda DNA binding sequence in a methylation-dependent manner (Fig. 6B). However, the Cada2^myxo^ showed Cada2-specific activity and could not induce *yfp* expression from the EcAda reporter (Fig. 6B). Together, these data illustrate the specificity, sufficiency and necessity of the identified regulatory features in the control of the adaptive responses across bacteria.

## Discussion

In this study, we identify Cada2, a novel methylation damage-specific transcription factor conserved across all major phyla of bacteria. We further implicate a critical and conserved role for PTM-based activation of this class of bacterial transcription factors, albeit with varying mechanisms of action (Cada2 vs EcAda) (Fig. 7A). We hypothesize that the organizational diversity observed across regulators of the adaptive response pathways are likely an adaptation to the physiological features of bacteria. The downstream adaptive response is, however, robust to this variability.

**Figure 7.**
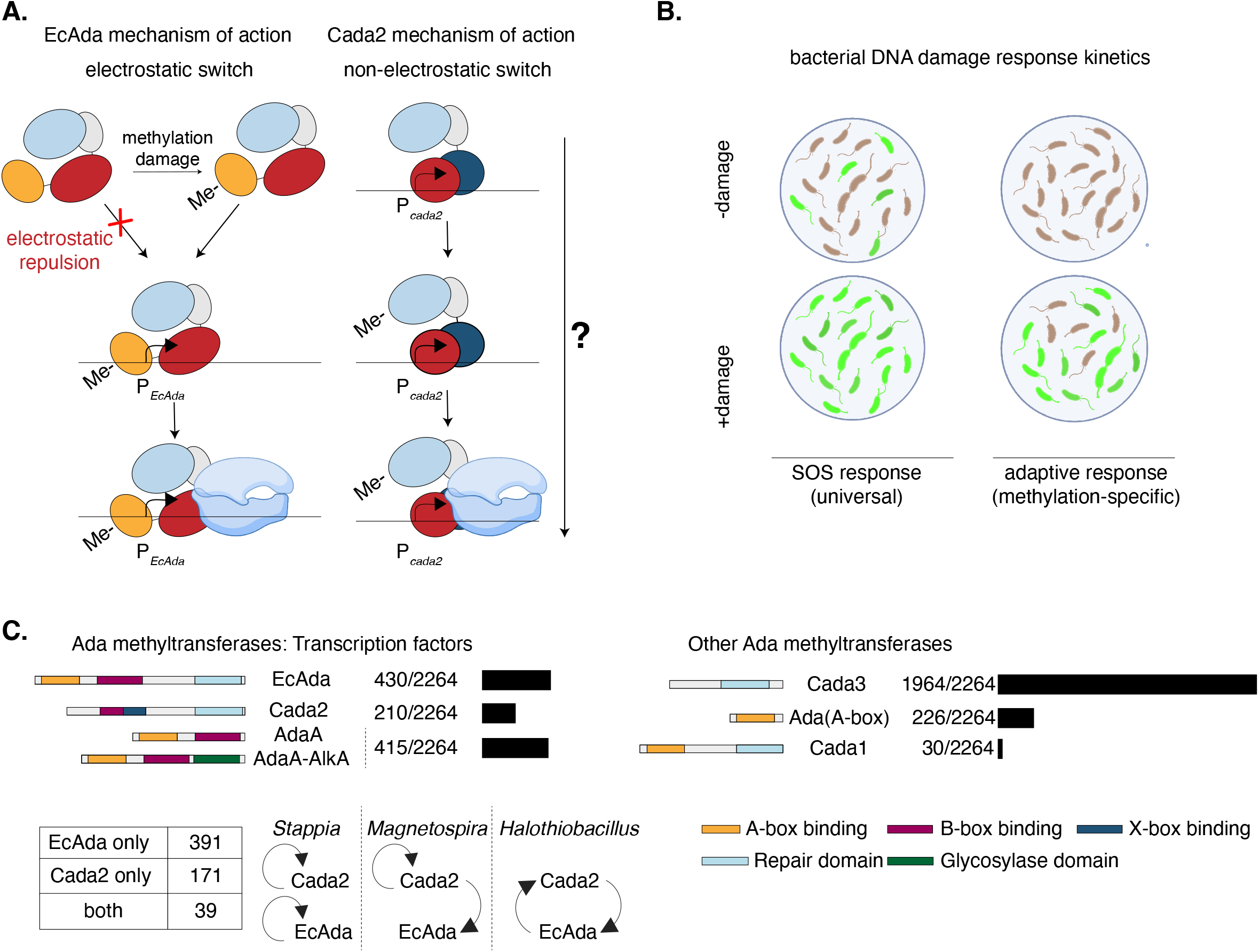
Diverse mechanisms, unifying functions of the bacterial adaptive response regulators. **(A)** EcAda and Cada2 are transcription factors for the bacterial adaptive response and function via two distinct mechanisms (see discussion section for details). **(B)** Although mechanistically distinct, features of the adaptive response between *E. coli* and *Caulobacter* are conserved (see discussion section for details). **(C)** [Top left] Domain organization of adaptive response regulatory methyltransferases and their prevalence estimated from a curated, non-redundant database of bacterial genomes at the species level. [Top right] Domain organization and prevalence similarly estimated for other adaptive response methyltransferases at the species level. [Bottom right] Functions associated with the PFAM domains that constitute these diverse organizations are annotated at the species level. [Bottom left] Table represents presence and absence of EcAda-like and Cada2-like proteins and their co-occurrence. In the instances of Cada2-EcAda co-occurrence, potential regulatory circuits predicted via identifying Cada2 and EcAda binding motif in their cognate promoters are shown.

### Diverse mechanisms, unifying function

The mechanistic differences between Cada2 and EcAda would suggest that the response dynamics would also be dissimilar. Cada2 and EcAda use distinct DNA binding domains to associate with their cognate promoters (B-box and ‘RLHD’ domains vs A and B-box binding domains respectively [26]. The two proteins are also activated as transcription factors in non-overlapping ways: In case of EcAda, methylation of the cysteine residue within the sequence-specific DNA-binding A-box domain activates it as a transcription factor [50,52]. In contrast, Cada2 activation occurs via PTM of the cysteine in its methyltransferase domain that is distally located from the sequence-specific DNA bindings domain. Furthermore, unlike EcAda, unmethylated Cada2 appears to form sequence-specific interactions with its own promoter (Fig. 3A, Fig. S2A). Presence of electrostatic repulsion (as in EcAda) would hinder this interaction. This would suggest that Cada2 acts via an alternate, non-electrostatic mechanism (Fig. 7A).

Yet despite these contrasts, the response kinetics are invariant between the two organisms (Fig. 7B). The adaptive response of *Caulobacter* is tightly regulated, showing low levels of expression (Ada OFF) in the absence of methylation DNA damage. Upon exposure to methylating agents, *Caulobacter* cells also elicit a delayed adaptive response, subsequent to the SOS response. Similar to the *E. coli* Ada response [22], the *Caulobacter* response exhibits bi-stability with a sub-population of cells resembling expression of Ada OFF cells even in the presence of DNA methylation damage. Interestingly, the OFF population is abolished in a strain over-expressing Cada2, mirroring observations made in case of EcAda [22] (Fig. 1B and Fig. 5A). The factors influencing the differences in the mechanisms of action of the response regulators, while retaining the defining features of the response itself are an important and exciting avenue for future investigations.

### Requirement for a methylation damage-specific response in bacteria

How pervasive is the adaptive response to methylation damage? Our computational analysis reveals that the regulatory Ada methyltransferase is widely conserved, but it appears to have diverse domain organizations (Fig. 7C). Given the apparent modularity of these domains (Fig. 7C), it is tempting to speculate that such diverse organizations arise as a result of domain-shuffling from a common ancestor protein [53–55]. Despite their prevalence, multiple regulators rarely occur on the same genome. A case in point is the comparison between EcAda and Cada2. We were able to detect only ∼7% genomes possessing both proteins. The genomes that encoded both EcAda and Cada2 displayed varied and unique regulatory circuits, including autoregulation as well as cross-regulation (Fig. 7C).

What drives the observed diversity? Comparison of EcAda and Cada2 might provide a compelling hypothesis. Both EcAda and Cada2 share the B-box motif in their cognate promoters. However, instead of the AT-rich A-box motif observed in the promoters of EcAda regulon genes, Cada2 binds to a GC-rich X-box motif. This correlates well with the GC-content of the genomes that encode these proteins. The GC-content of the *E. coli* genome is ∼50% (low), while the GC-content of the *Caulobacter* genome is ∼67% (high). This hypothesis is consistent with previous studies that have also suggested the interdependence of genome GC-content and presence/absence of specific DNA repair pathways [56–59].

Thus, it is likely that distinguishing physiological characteristics of bacteria warrant organizational adaptation and domain reorganizations of regulatory Ada methyltransferases. However, regardless of the diversity observed at a regulator level, the adaptive response-like pathways seem to retain certain characteristics as emphasized previously. In this context, we highlight the PTM-based transcriptional switch required for activation and regulation of the response. It is possible that costs (e.g. Fig. 5D) or fitness advantages associated with this response serve as constraining forces that render this system robust to variations in regulator organization, and yet drive the retention of key regulatory features. Together, the conservation of bacterial adaptive response mechanisms, albeit in diverse regulatory forms, underscores the fundamental requirement for a dedicated methylation-specific DNA damage response.

## Materials and Methods

### Bacterial strains and growth conditions

All strains, plasmids and oligos used in this study are listed in Supplemental Table S1-S3 respectively. *Caulobacter crescentus* cells were grown at 30°C in Peptone Yeast Extract (PYE) media (0.2% peptone, 0.1% yeast extract and 0.06% MgSO_4_) and supplemented with antibiotics or inducers, as required, at suitable concentrations. For induction of *cada2* expression, 0.3% xylose was introduced into the cultures, unless otherwise stated.

### Fluorescence microscopy and image analysis

Imaging was performed on an epifluorescence, wide-field microscope (Eclipse Ti-2E, Nikon) with a 60x/1.4NA oil immersion objective and a motorized stage. pE-4000 (CoolLED) was used as the LED excitation source. Exposure times for all *yfp* samples (λ=490nm) was 500ms and exposure was maintained at 50% of LED power. Images were captured using a Hamamatsu Orca Flash 4.0 camera. Focus was ensured via an infrared-based Perfect Focusing System (Nikon). Samples were prepared as detailed in [60]. For time course microscopy, 1mL of cultures were aliquoted at various time points, pelleted and resuspended in appropriate volume of growth media. 2µL of the resuspended cells were spotted on 1% agarose (Invitrogen ultrapure) pads and imaged. For time lapse microscopy, 2µL of culture were spotted on 1.5% low-melting GTG agarose pads supplemented with PYE media and 1.5mM MMS damage. Throughout the duration of the time lapse, samples were grown in an OkoLab incubation chamber maintained at 30°C and imaged at regular intervals. Cell segmentation and fluorescence intensity analysis was carried out via Oufti software [61].

### RNA-sequencing

#### Sample collection

Overnight cultures of *Caulobacter* cells were back-diluted to 0.025 OD_600_ and grown at 30°C for 3 hours. When OD_600_ of culture was ∼0.1, DNA damage (MMS (1.5mM) / MMC (0.25 μg/ml) / norfloxacin (8 μg/ml)) was introduced. 2 mL of *Caulobacter* cells were harvested and spun at 10000g for 5 minutes. Supernatant was then discarded and the cell pellets were snap frozen with liquid nitrogen and stored at –80°C until the samples were further processed. Cells were harvested at 0, 20 and 40 minutes post DNA damage exposure.

#### RNA-extraction

RNA was extracted from the cell pellets according to a previous study [49]. Cells were lysed using 400 μL of pre-heated Trizol in a thermomixer at 65°C, 2000rpm. The lysed cells were transferred to –80°C for a minimum of 30 minutes and were then centrifuged at 4°C. The supernatant was carefully transferred to 100% ethanol of equivalent volume. RNA was then extracted via the protocol mentioned in the Direct-zol RNA MiniPrep (Zymo, Cat. no. R2052) kit. The extracted RNA was then subjected to DNAse treatment in order to remove any genomic DNA from the sample. Total RNA from the samples was purified using the RNA Clean & Concentrator-25 (Zymo, Cat. no. R1018) kit. Integrity of the RNA was tested using Bioanalyzer instrument. mRNA was isolated from the total sample using the RiboMinus kit (Thermo, Cat. no. K155004) and submitted for RNA-seq at the NCBS next generation sequencing facility.

#### Library preparation and RNA-seq

Kits used for preparing RNA libraries and sequencing platform used for RNA-seq are listed below.

| background | treatment condition | RNA library preparation kit | Sequencing platform |
| --- | --- | --- | --- |
| *wild type* | MMS | NEBNext® Ultra™ II Directional RNA Library Prep with Sample Purification Beads (Catalog no-E7765L) | Illumina Hiseq2500 |
| *ΔrecA* | MMS |  |  |
| *Δcada1* | MMS |  |  |
| *Δcada2* | MMS |  |  |
| *wild type* | MMC | NEBNext® Ultra™ II Directional RNA Library Prep with Sample Purification Beads (Catalog no-E7765L) | Illumin Novaseq4000 |
| *wild type* | norfloxacin |  |  |

##### RNA-sequencing analysis

Raw reads for the sequencing results were obtained as fastq files. The reference genome sequence (.fna) and annotation (.gff) files for the *Caulobacter crescentus NA1000* strain (accession number: <u>NC_011916.1</u>) were downloaded from the NCBI file transfer protocol (ftp) website (“ftp.ncbi.nlm.nih.gov”). The raw read quality was checked using the FastQC software (version 0.11.5). Burrows-Wheeler Aligner (BWA) (version 0.7.17-r1188) was used to index the reference genome. Reads with raw read quality >= 20 were aligned to the indexed genome using the BWA aln -q option. Samtools (version 0.1.7) was used to filter out multi-mapped reads (). Bedtools (version 2.26.0) was used to calculate the reads count per gene using the annotation file (.bed) [62]. The normalization and differential gene expression analysis for the conditions were carried out using edgeR [63]. To estimate differential gene expression under DNA damaging agent and for the deletion mutants zero hour values for each respective samples were used for comparison. All samples with deletion mutants were compared with their respective 0 hour samples. For *wild type*, the 4 replicates of 0 hour samples were grouped and compared to other *wild type* samples to calculate differential gene expression under MMC, MMS and norfloxacin. A gene was defined as up-regulated based on two thresholds; in a certain condition if the log_2_ fold change (from control) of the gene is greater than or equal to 1 (i.e. the gene content has doubled in this condition) and if the false discovery rate across replicates was less than 0.05. Similarly, a gene was down-regulated in that condition if the gene’s log_2_ fold change from control was less than -1 and the false discovery rate across replicates is less than 0.05. DGE analysis was done using Google Colab with R version 4.2.3 (2023-03-15).

##### Western blotting

Western blotting was performed as described in a previous study [64]. Briefly, overnight cultures in PYE media was back-diluted to 0.025 OD_600_ and grown at 30°C for 3 hrs. At ∼0.1OD_600_, 1.5mM MMS was added to the culture. 0, 2 and 4 hours post introduction of DNA damage, cells corresponding to 0.6 OD_600_ were harvested. The culture was pelleted following which supernatant was discarded. The pelleted cells were resuspended in 200 μL of lysis buffer [500mM Tris-HCL (pH6.8), 8% SDS, 40% glycreol, 2-Mercaptoethanol and sufficient amount of bromophenol blue] and heated at 95°C for 3 minutes. This was followed by two rounds of 30 second vortex spins, each followed by short spins. The resuspended cells were again heated to 95°C post which samples were loaded on to a 10% SDS-PAGE gel and subjected to electrophoresis. The resolved protein bands were then transferred on to a PVDF membrane and probed with anti-flag antibodies for estimation of *cada2* levels. The same samples were also probed with anti-RpoA antibodies in order to verify uniform loading of samples. The blots were developed using SuperSignal West PICO PLUS chemiluminescent substrate.

#### Mass spectrometry

##### Sample collection

Cells over-expressing flag-tagged *wild type cada2* or its mutants (*cada2_R68A_*, *cada2_R114A_*) from a xylose-inducible promoter in a *Δcada2* background were used. Overnight cultures grown in PYE media containing gentamycin was back-diluted to 0.025 OD_600_ and grown at 30°C for 3 hrs. At ∼0.1OD_600_, 1.5mM MMS and 0.3% xylose was added to the culture and culture volume corresponding to 0.6 OD_600_ was collected after 4 hrs of treatment. The culture was pelleted following which supernatant was discarded. The pelleted cells were resuspended in 200 μL of lysis buffer [500mM Tris-HCL(pH6.8), 8% SDS, 40% glycreol, 2-Mercaptoethanol and sufficient amount of bromophenol blue] and heated at 95°C for 3 minutes. This was followed by 30 second vortex spin, followed by a short spin. The above two steps were repeated twice. The resuspended cells were again heated to 95°C post which samples were loaded on to a 10% SDS-PAGE gel and subjected to electrophoresis. The gel was appropriately resolved and stained with Coomassie dye. Bands corresponding to the size of Cada2-3x-Flag (∼37 kDa) were excised from the gel, resuspended in milliQ water and submitted to the NCBS mass spectrometry facility for further processing.

##### Sample preparation and mass spectrometry

The excised gel fragments were first washed thrice with LC-MS water post which the supernatant water was discarded. Next, the gel pieces were chopped followed by destaining with 1:1 100 mM Triethyl ammonium bicarbonate (TEAB) with 100% acetonitrile (ACN). Post destaining, the supernatant was discarded. ∼200 μL of 100% ACN was then added and the sample was kept at room temperature till the gel pieces shrunk, following which the supernatant was discarded. The previous step of ACN addition was repeated for a second time. The gel pieces were allowed to dry for a few minutes at room temperature. Next, the gel was suspended in a minimum of 500 ng of Trypsin with 100-200 μL of 100mM TEAB. The samples were then kept for overnight trypsin digestion at 37 °C. 100 μL of 100% ACN with 0.1% Formic acid was then introduced to the digested sample and the digested sample were sonicated for 5 min. The supernatant was then collected into a fresh tube. The post-digestion step was repeated for a second time. The collected supernatant was then dried using a SpeedVac vaccum concentrator and reconstituted in 0.1% formic acid. This was followed by desalting post which samples were injected into Orbitrap Fusion Tribrid Mass Spectrometer coupled to a Thermo EASY nanoLC 1200 chromatographic system for mass spectrometry. The results were analyzed using the PeakStudio 8.0 for identifying peptide fragments and their associated post-translational modifications (if any).

#### Chromatin Immunoprecipitation Sequencing (ChIP-Seq)

Fifty mL PYE was sub-inoculated with overnight *Caulobacter crescentus* cultures to OD_600_ of 0.025, cultures were grown at 30°C for another 3 hrs with shaking at 250rpm until reaching OD_600_ of 0.1. MMS (final concentration of 1.5 mM) was then added, and the cultures were outgrowth for another 2 or 4 hrs before fixation with formaldehyde. Controls of no MMS treatment were also included. Cells were fixed with formaldehyde (final concentration of 1%) at room temperature for 30 min, then quenched with 0.125 M glycine for another 15 min at room temperature. Cells were washed three times with 1x PBS (pH 7.4) and resuspended in 1 mL of lysis buffer [20 mM K-HEPES pH 7.9, 50 mM KCl, 10% glycerol, and Roche EDTA-free protease inhibitors]. ChIP-seq was performed as described previously in Tran *et al* (2018) [65]. Briefly, the cell suspension was sonicated on ice using a Soniprep 150 probe-type sonicator (11 cycles, 15s on, 15s off, at setting 8) to shear the chromatin to below 1 kb, and the cell debris was cleared by centrifugation (20 min at 13,000 rpm at 4°C). The supernatant was then transferred to a new 2 mL tube and the buffer conditions were adjusted to 10 mM Tris-HCl pH 8, 150 mM NaCl and 0.1% NP-40. Fifty microliters of the supernatant were transferred to a separate tube for control (the input fraction) and stored at -20°C. Meanwhile, antibodies-coupled beads were washed off storage buffers before being added to the above supernatant. α-M2 FLAG antibodies coupled to sepharose beads (Merck, UK) were employed for ChIP-seq of Cada2-FLAG and RpoC-FLAG. Briefly, 100 μL α-FLAG beads were washed off the storage buffer by repeated centrifugation and resuspension in IPP150 buffer [10 mM Tris-HCl pH 8, 150 mM NaCl and 0.1% NP-40]. Beads were then introduced to the cleared supernatant and incubated with gentle shaking at 4°C overnight. Beads were then washed five times at 4°C for 2 min each with 1 mL of IPP150 buffer, then twice at 4°C for 2 min each in 1x TE buffer [10 mM Tris-HCl pH 8 and 1 mM EDTA]. Protein-DNA complexes were then eluted twice from the beads by incubating the beads first with 150 μL of the elution buffer [50 mM Tris-HCl pH 8, 10 mM EDTA, and 1% SDS] at 65°C for 15 min, then with 100 μL of 1X TE buffer + 1% SDS for another 15 min at 65°C. The supernatant (the ChIP fraction) was then separated from the beads and further incubated at 65°C overnight to completely reverse the crosslink. The Input fraction was also de-crosslinked by incubation with 200 μL of 1X TE buffer + 1% SDS at 65°C overnight. DNA from the ChIP and input fraction were then purified using the PCR purification kit (Qiagen) according to the manufacturer’s instruction, then eluted out in 40 µL water. Purified DNA was then made into libraries suitable for Illumina sequencing using the NEXT UltraII library preparation kit (NEB, UK). ChIP libraries were sequenced on the Illumina Hiseq 2500 at the Tufts University Genomics facility.

#### ChIP-seq analysis

Raw reads for the ChIP-seq results were obtained as fastq files. BWA was used in order to align the reads to the *Caulobacter* genome similar to analysis of RNA-seq data. The aligned reads in .bam format were converted into a .bed file format using the bamtobed option of Bedtools. Coverage at each nucleotide position was calculated using the Bedtools genomecov command. Subsequently, ChIP-seq profiles from these coverage files were plotted using a custom python script. Pearson correlation between ChIP-seq profiles for *cada2 flag* and *rpoC flag* was calculated using Numpy. Peakzilla was used in order to identify bona fide enrichment peaks. .bed files from *cada2 3xflag* or rpoC 3xflag ChIP-seq under MMS exposure were compared to untagged *wild type* cells treated under identical conditions for peak-calling.

#### ChIP and Quantitative PCR analysis

For quantitative-PCR (q-PCR) analysis of *cada1* and *cada3* promoter enrichment in ChIP experiments, purified DNA from both input and ChIP fractions was diluted 1:4 in water and 1 μL was used for qPCR using a SYBR^®^ Green JumpStart^™^ *Taq* ReadyMix^™^ (CAT S4438, Merck, UK) and a BioRad CFX96 instrument. Fold enrichments were calculated using the comparative Ct method (ΔΔCt) and represent the relative abundance of *cada1* and *cada3* promoter DNA compared to *rpoD* DNA as a negative control. All fold enrichment values represent the average of three biological replicates.

The following oligos were used in qPCR reactions:

cada1 (fw, TAGGACGCGACTGCTGA, rv, TCCTTTCGTGAGGAGACCA)

cada3 (fw, ATCGCCCGCATGGAATAC, rv, AGAAGGAAGCTACTACCGGAT)

rpoD (fw, TCAGGCCAAGAAGGAAATGG, rv, GCCTTCATCAGGCCGATATT)

#### Survival assay

Overnight cultures of *Caulobacter* strains were back-diluted to OD_600_ 0.1. The cultures were incubated at 30°C for 3 hours. All cultures were then normalized to 0.3 OD_600_ and serially diluted in ten-fold increments (10^-1^ -10^-8^). 6 μl of each dilution was spotted on PYE plates containing appropriate chemicals (DNA damaging agents/antibiotics/inducer). The plates were incubated at 30°C for 48 hours post which survival of the respective strains was determined by counting the number of spots.

#### Computational predictions of protein structures via AlphaFold

Three-dimensional structural predictions for EcAda, Cada1, Cada2 and Cada3 were made by ColabFold (AlphaFold2 coupled with MMSeq2 hosted on Google Colaboratory) [47,48]. Primary sequence of each protein was used as query sequence. Default parameters were used for other settings. Structural models shown in this paper are models with the highest pLDDT score.

#### Computational analysis of protein conservation

All “complete” and “latest” (assembly_summary.txt; as of January 2017) genome information files for ∼6000 bacteria were downloaded from the NCBI ftp website using in-house scripts for whole genome sequences (.fna), protein coding nucleotide sequences (.fna), RNA sequences (.fna) and protein sequences (.faa). All the organisms were assigned respective phylum based on the KEGG classification. (https://www.genome.jp/kegg/genome.html; as of May 2018).

For identification of Ada domains and Ada variants, initial blastp was run using each of the four protein domain sequences of EcAda from *Escherichia coli MG1655* as the query sequence against the UniprotKB database with an E-value cutoff of 0.0001. The four domains of Ada include A-box, B-box, RNAse-like and repair. The top 1000 full-length sequence hits were downloaded from UniProt for all the four domains. A domain multiple sequence alignment (MSA) was made using phmmer –A option with the top 1000 hits as the sequence database and *E. coli* domain sequences as the query. An hmm profile was built using the hmmbuild command for the MSA obtained in the previous step. To find domain homologs, hmmsearch command with an E-value cutoff of 0.0001 was used with the hmm profile as the query against a database of 5,973 bacterial genome sequences. These homolog searches were done using HMMER package v3.3. Different Ada variants were identified based on the assignment of the different combinations of Ada domains to the same protein.

A phylogenetic species tree was constructed using 16S rRNA sequences. One sequence per genome was extracted from a multi-fasta file. An MSA was built using muscle v3.8.31 with default options, followed by alignment trimming using BMGE v1.12. Using IQTREE v1.6.5, a maximum likelihood-based phylogeny was built with the best model chosen using ModelFinder (-m MF option) against 285 other models. Branch supports were assessed using both 1000 ultrafast bootstrap approximations (-bb 1000 –bnni option) and SH-like approximate likelihood ratio test (- alrt 1000 option). Final tree for visualization was pruned to contain four major phyla – Proteobacteria, Actinobacteria, Firmicutes and Bacteroidetes. Online tool Itol [66] was used for visualization and EcAda and Cada2 presence/absence was overlaid on the phylogeny.

For identification of essential regulatory motifs of Cada2 and EcAda, protein blast was run on the coding sequences of EcAda or Cada2 against the UniprotKB database with an E-value cutoff of 10. The top 1000 full-length sequence hits were downloaded from UniProt for both proteins and were submitted to the MEME discovery program in order to identify conserved features.

## Author contributions

AK, TL and AB conceived the project. AK led the project, carried out majority of the experiments, generated tools and reagents, and conducted data analysis. NT and TL carried out ChIP-seq and ChIP-qPCR experiments. MS contributed to bioinformatic analysis. NS contributed to RNA-seq analysis. AB procured funding and supervised the project. AK and AB wrote the manuscript, with inputs from all authors.

## Acknowledgements

We thank Dr. Asha Joseph for contribution to RNA-sequencing experiments and Kaustav Mitra for carrying out initial bacterial-two-hybrid experiments. We are grateful to members of the AB lab for helpful comments and discussions. We acknowledge support from NCBS Mass Spectrometry facility and the Next Generations Sequencing facility for key experiments. This work was supported by funding to AK (TIFR graduate student fellowship) and to AB via the India Alliance Intermediate Grant (Grant number IA/I/21/1/505630) and intramural funding via NCBS-TIFR (Grant number 03/3/2019/R&D-II/DAE/4749). This work was also supported by the Lister Institute fellowship, the Wellcome Trust Investigator grant 221776/Z/2/Z (to T.B.K.L), and by the Biotechnology and Biological Sciences Research Council grant-in-aid (BBS/E/J/000PR9791 that supports N.T.T).

## Data availability

All sequencing data are deposited in GEO, with accession number GEO.

## Supplementary Information

### Supplementary figure legends

**Figure S1.**
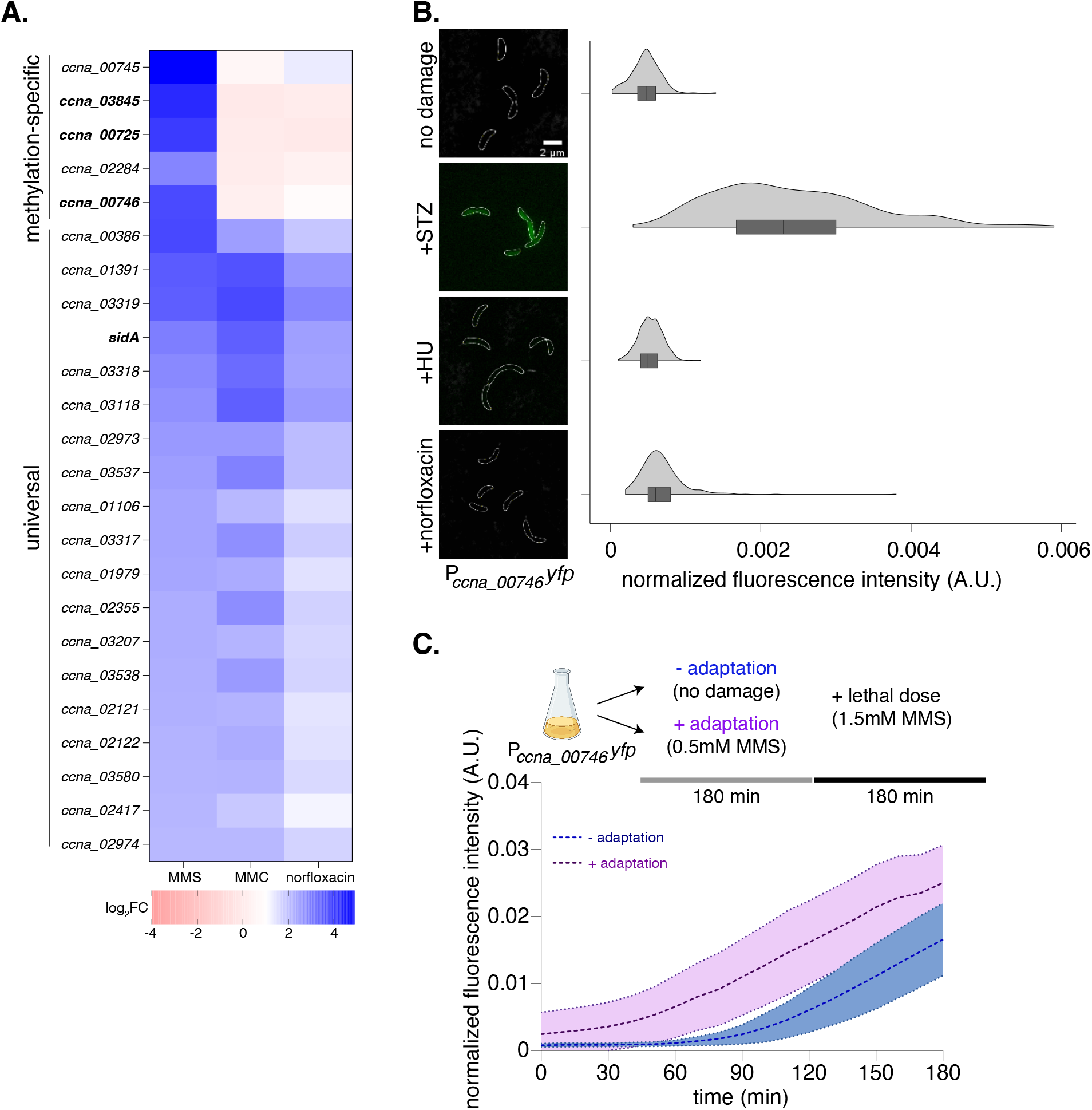
A methylation-specific DNA damage response in Caulobacter. **(A)** Heat map of log_2_FC values for genes upregulated in wild type cells following MMS, MMC or Norfloxacin treatment. **(B)** [Left] Representative cells showing P*_ccna_00746_-yfp* reporter induction upon exposure to STZ, MMC, norfloxacin and HU. [Right] Violin plots show fluorescence intensity distribution normalized to cell area from single cells (n=300, from three biological replicates). **(C)** [Top] Schematic of the experimental protocol for testing the adaptive property of the *Caulobacter* methylation-specific damage response. Cultures of P*_ccna_00746_-yfp* cells were exposed to a sub-lethal dose of 0.5mM either MMS (adapted) or no MMS (non-adapted). The cells were subsequently exposed to a higher dose of 1.5mM MMS on agarose pads supplemented with PYE medium. [Bottom] Normalized fluorescence intensity kinetics was measured via time lapse microscopy over 3 hours of lethal MMS exposure. Dotted lines (in dark) indicate mean time trace of induction kinetics while the shaded region (in light) indicates the standard deviation of all time traces for the respective conditions (here and for all other time lapse data) (n=25).

**Figure S2.**
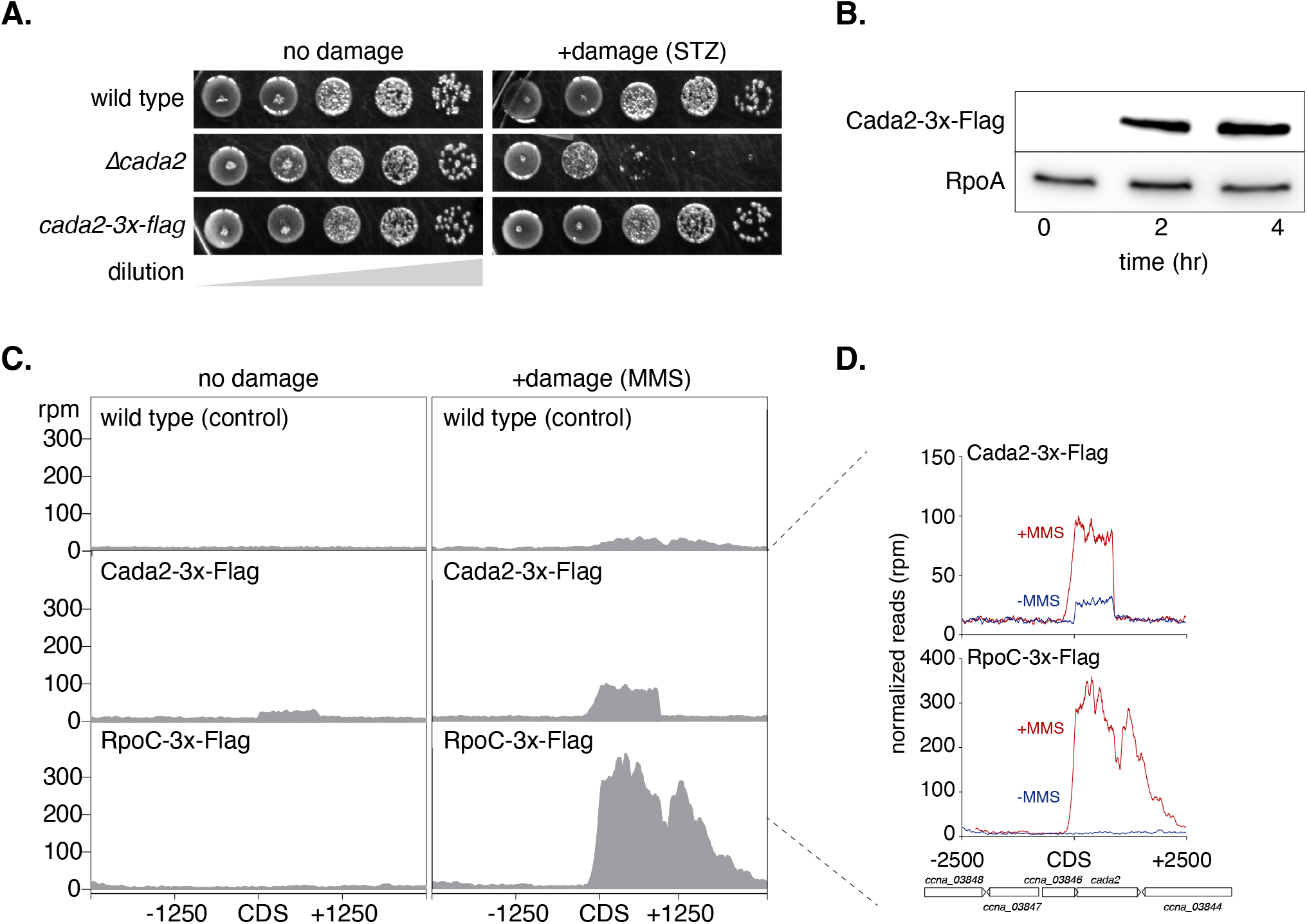
Cada2 associates with adaptive response promoters in a sequence-specific manner. **(A)** Survival assay of *cada2-3x-flag* strain in the presence or absence of STZ damage (5μg/ml). **(B)** Western blot showing Cada2-3x-flag levels at 0, 2 and 4 hours after 1.5mM MMS exposure. As a loading control, RpoA is probed. **(C)** ChIP-seq profiles for wild type (control), Cada2-3x-Flag or RpoC-3x-Flag ±2.5kb around *cada2* CDS before (no damage) and after (+damage) exposure to 1.5mM MMS. **(D)** Zoomed-in ChIP-seq profiles for Cada2-3x-Flag and RpoC-3x-Flag ±2.5kb around *cada2* CDS before and after exposure to 1.5mM MMS.

**Figure S3.**
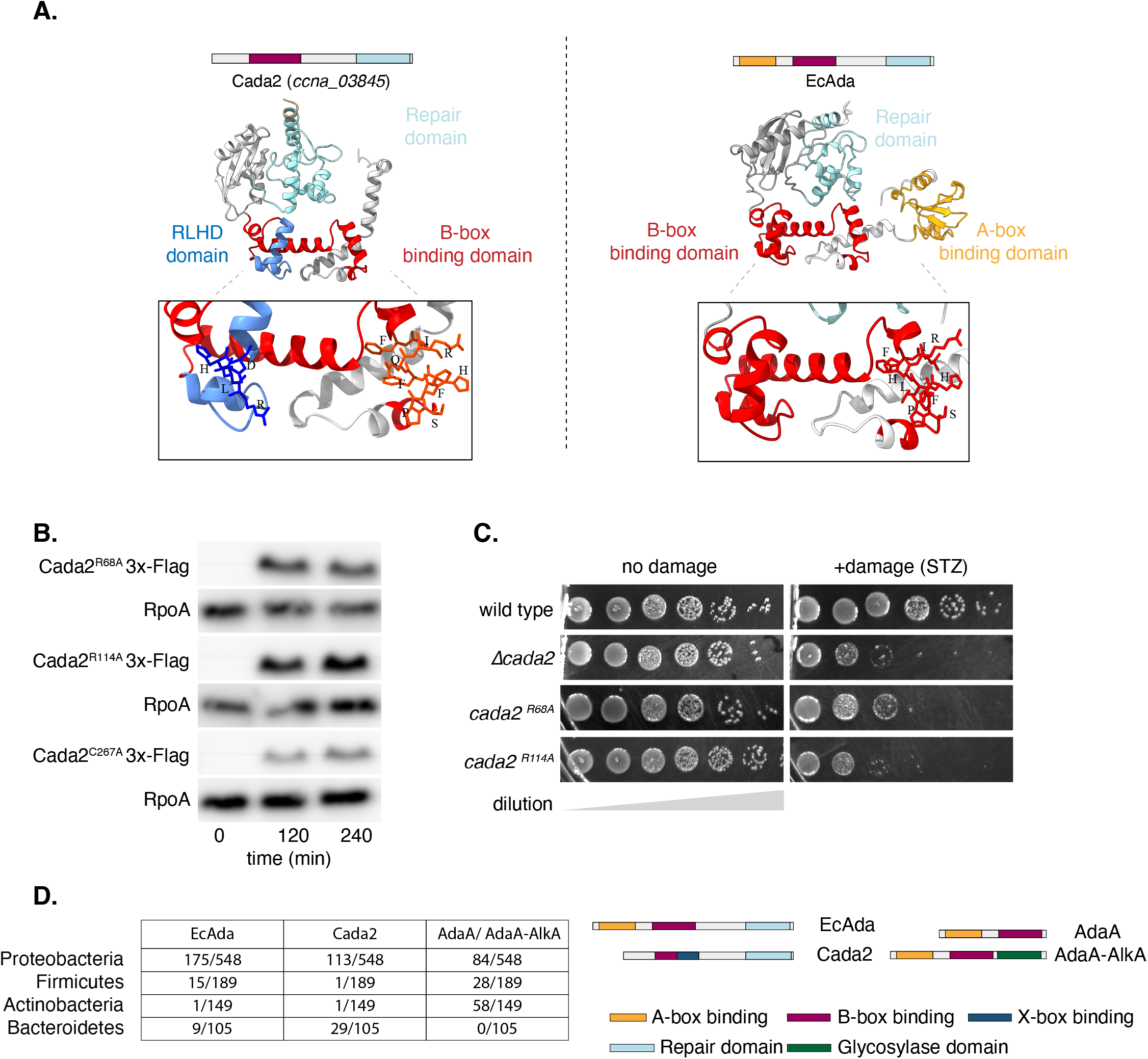
Cada2-like proteins are widespread and encode a novel DNA binding domain. **(A)** (Top) Domain organization and predicted Alphafold structure of Cada2 and EcAda is indicated. (Bottom) Closeup of the EcAda and Cada2 regulatory domain reveals that the Cada2 B-box binding domain (possessing the ‘SPFHQR’ amino acid sequence) and the newly identified sequence-specific binding domain of Cada2 (possessing the ‘RLHD’ amino acid sequence) are part of a helix-turn-helix domain similar to the B-box binding domain of EcAda. The position of these conserved motifs are highlighted and labelled as a ball-and-stick in the overall ribbon representation of the models. **(B)** Western blot of flag-tagged Cada2 mutants (Cada2^R68A^ and Cada2^R114A^) overexpressed in a *Δcada2* strain from a xylose-inducible promoter treated with MMS damage. **(C)** Survival assay of *cada2^R68A^ and cada2^R114A^* with or without STZ damage. **(D)** [Left] Prevalence of EcAda-like, Cada2-like and AdaA-like proteins estimated from a curated, non-redundant database of bacterial genomes is analysed at the genus level. Numbers represent the presence of these proteins for the major bacterial clades. [Right] domain organization of the respective adaptive response regulatory proteins analysed here.

**Figure S4.**
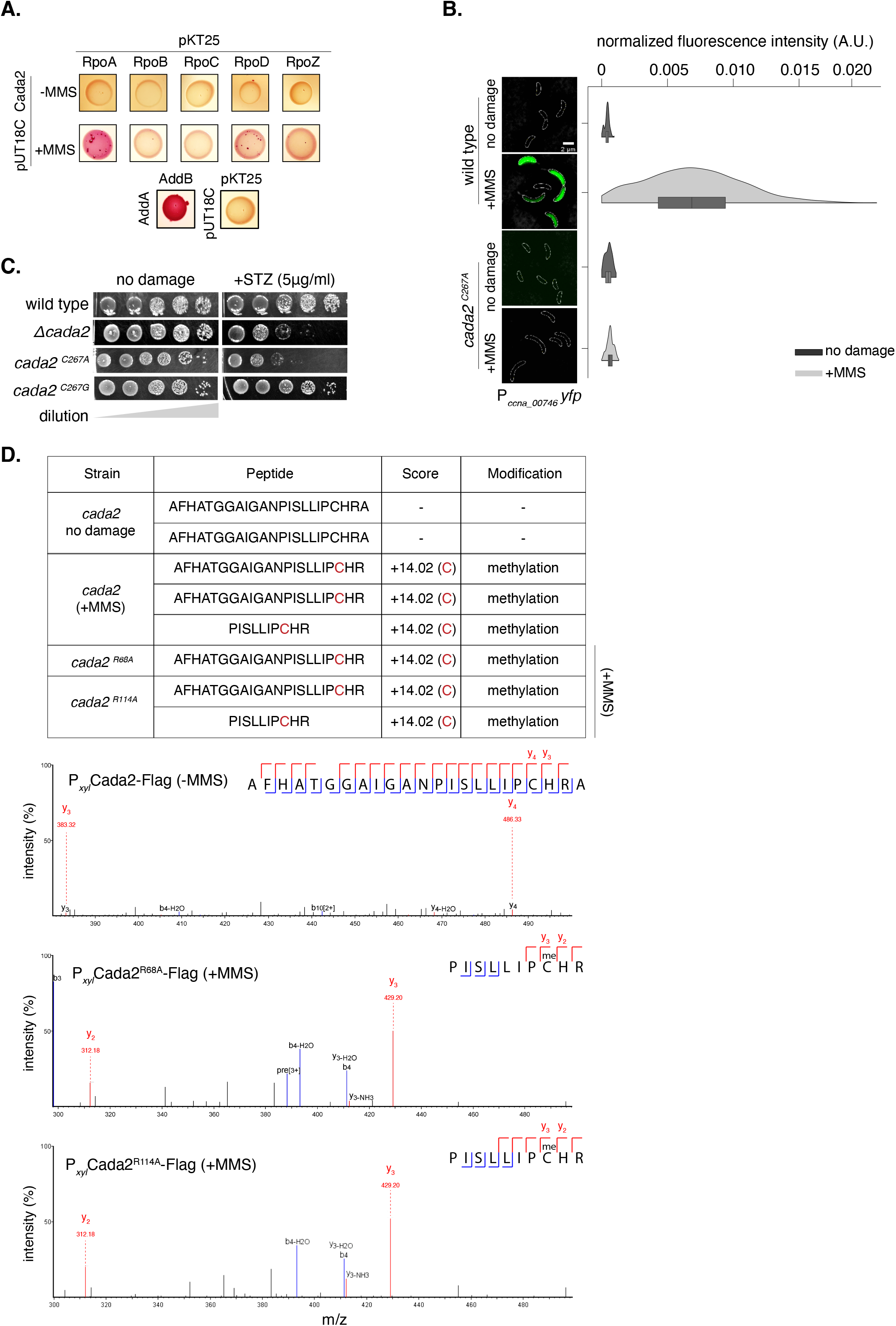
Cada2 is a methylation-dependent transcription factor. **(A)** Bacterial two-hybrid assay to test interaction between Cada2 and RNA polymerase subunits. T18-Cada2 was tested for interaction with T25-RNA polymerase holoenzyme subunits with and without 1.5mM MMS. The presence of red colonies indicated positive interaction. As a positive control AddA (T18) and AddB (T25) are used, and empty vectors (T18 and T25) are used as negative control. Representative images from 2 independent repeats are shown. **(B)** [Left] Representative cells showing P*_ccna_00746_-yfp* reporter induction in wild type (from Fig. 1B) and c*ada2^267A^* mutant background under 1.5mM MMS damage. [Right] Violin plots showing fluorescence intensity distribution normalized to cell area from single cells in the presence (dark) and absence of damage (light) (n=200, from two biological replicates). **(C)** Survival assay of the *cada2 mutants* (*cada2^C267A^* or c*ada2^C267G^*) with and without STZ damage (5μg/ml). (D) [Above] Table representing peptides corresponding to the Cada2 methyltransferase domain bearing the ‘PCHR’ motif as identified via mass spectrometry. In the absence of MMS, peptides corresponding to wild type Cada2-flag are unmethylated. Upon exposure to MMS damage, peptides corresponding to wild type as well as mutant Cada2-3x-Flag exhibit methylation at the Cys^267^ residue. [Below] Representative mass spectrometry fragmentation patterns for peptides in the above table. In case of wild type (no damage) representative spectrum from 3 fragments across two biological replicates is shown. No methylation modification on Cys^267^ was detected. In case of Cada2^R68A^ representative spectrum from 4 fragments across two biological replicates is shown. Methylation modification on Cys^267^ was detected in 3 out of 4 fragments. In case of Cada2^R114A^ representative spectrum from 7 fragments across two biological replicates is shown. Methylation modification on Cys^267^ was detected in 5 out of 7 fragments.

**Figure S5.**
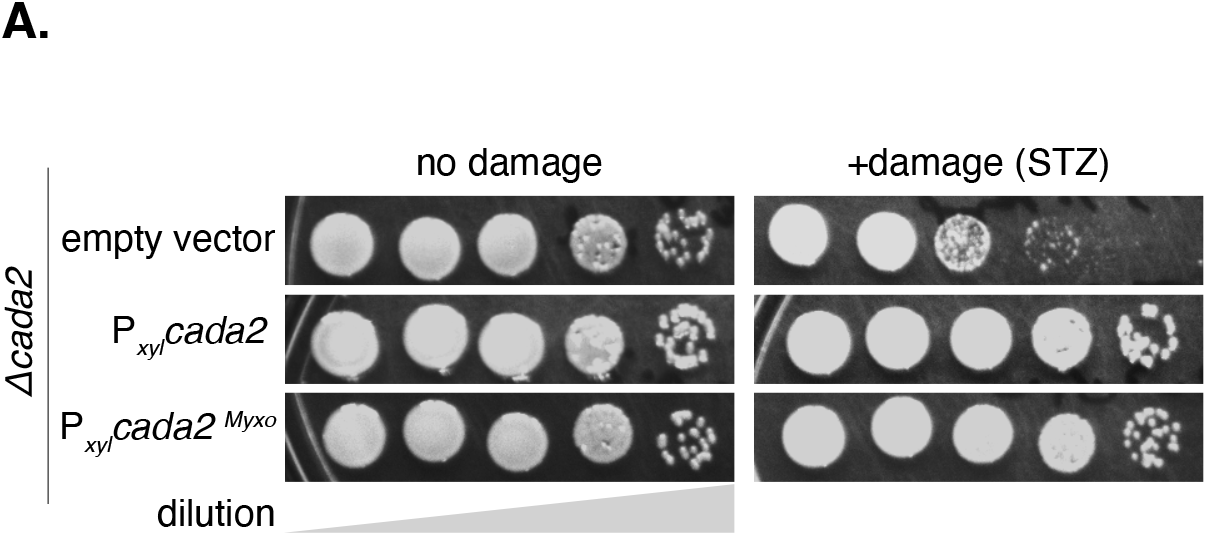
Regulatory features of Cada2 are conserved across bacteria and distinct from the EcAda paradigm. **(A)** Survival assay of *Δcada2* strain overexpressing *cada2^caulo^ or cada2^myxo^* (from xylose-inducible promoter) under methylation damage (5μg/ml STZ). Survival of these strains was compared to a control strain comprising of an empty vector in a *Δcada2* background.

**Table S1:** Strains used in present study.

| Strain | Genotype | Strain construction |
| --- | --- | --- |
| CB15N | NA1000 | NA |
| NABC2 | *CB15N*; $\Delta recA$ | [1] |
| NABC581 | CB15N; $P_{sidA}$-YFP::kan<sup>R</sup> | [2] |
| NABC477 | BTH101; pKT25-addB::kanR ; pUT18C-addA::carbR | [3] |
| NABC741 | CB15N; *rpoC*-flag::spec<sup>R</sup> | [4] |
| NABC735 | BTH101; pKT25; pUT18C | [5] |
| NABC706 | CB15N; $P_{ccna\_00746}$-YFP::kan<sup>R</sup> | CB15N strain was transformed with pNABC735 plasmid to integrate $P_{ccna\_00746}$-YFP linked to kan<sup>R</sup> under the $P_{xyl}$ locus. |
| NABC707 | *CB15N*; $\Delta cada1$ | CB15N was transformed with pNABC736 plasmid to generate deletion of *cada1* through two-step recombination procedure (Skerker et al., 2005). |
| NABC708 | *CB15N*; $\Delta cada2$ | CB15N was transformed with pNABC737 plasmid to generate deletion of *cada2* through two-step recombination procedure (Skerker et al., 2005). |
| NABC709 | *CB15N*; $\Delta cada3$ | CB15N was transformed with pNABC738 plasmid to generate deletion of *cada3* through two-step recombination procedure (Skerker et al., 2005). |
| NABC710 | *CB15N*; $\Delta cada1$; $P_{ccna\_00746}$-YFP::kan<sup>R</sup> | NABC707 strain was transformed with pNABC735 plasmid to integrate $P_{ccna\_00746}$-YFP at the *xyl* locus. |
| NABC711 | *CB15N*; $\Delta cada2$; $P_{ccna\_00746}$-YFP::kan<sup>R</sup> | NABC708 strain was transformed with pNABC735 plasmid to integrate $P_{ccna\_00746}$-YFP at the *xyl* locus. |
| NABC712 | *CB15N*; $\Delta cada3$; $P_{ccna\_00746}$-YFP::kan<sup>R</sup> | NABC709 strain was transformed with pNABC735 plasmid to integrate $P_{ccna\_00746}$-YFP at the *xyl* locus. |
| NABC713 | *CB15N*; $\Delta cada2$; pMT687-$P_{xyl}$-*cada2*::gent<sup>R</sup> | NABC708 was transformed with pNABC739 plasmid. |
| NABC714 | CB15N; *cada2* (C267A) | CB15N was transformed with pNABC740 plasmid to generate a specific mutation in the conserved C267 residue of *cada2* through two-step recombination procedure (Skerker et al., 2005). |
| NABC715 | CB15N; *cada2* (C267A); $P_{ccna\_00746}$-YFP::kan<sup>R</sup> | NABC714 strain was transformed with pNABC735 plasmid to integrate $P_{ccna\_00746}$-YFP at the *xyl* locus. |
| NABC716 | CB15N; *cada2* (C267G) | CB15N was transformed with pNABC741 plasmid to generate deletion of *cada1* through two-step recombination procedure (Skerker et al., 2005). |
| NABC717 | CB15N; *cada2* (C267G); $P_{ccna\_00746}$-YFP::kan<sup>R</sup> | NABC716 strain was transformed with pNABC735 plasmid to integrate $P_{ccna\_00746}$-YFP at the *xyl* locus. |
| NABC718 | CB15N; *cada2*-flag::spec<sup>R</sup> | CB15N strain was transformed with pNABC742 plasmid to integrate *cada2*-flag linked to spec<sup>R</sup> under the endogenous *cada2* locus. |
| NABC719 | CB15N; $P_{ccna\_00746}$-*YFP(scramble)::kan<sup>R</sup>* | CB15N strain was transformed with pNABC743 plasmid to integrate $P_{ccna\_00746}$-*YFP(scramble)* linked to *kan<sup>R</sup>* under the $P_{xyl}$ locus. |
| NABC720 | CB15N; $P_{ccna\_00746}$-*YFP(AT-rich)::kan<sup>R</sup>* | CB15N strain was transformed with pNABC744 plasmid to integrate $P_{ccna\_00746}$-*YFP(AT-rich)* linked to *kan<sup>R</sup>* under the $P_{xyl}$ locus. |
| NABC721 | CB15N; *cada2* (R68A) | CB15N was transformed with *cada2* pNABC745 plasmid to generate a specific mutation in the conserved R68 residue of *cada2* through two-step recombination procedure (Skerker et al., 2005). |
| NABC722 | CB15N; *cada2* (R68A); $P_{ccna\_00746}$-*YFP::kan<sup>R</sup>* | NABC721 strain was transformed with pNABC735 plasmid to integrate $P_{ccna\_00746}$-YFP at the *xyl* locus. |
| NABC723 | CB15N; *cada2* (R114A) | CB15N was transformed with pNABC746 plasmid to generate a specific mutation in the conserved R114 residue of *cada2* through two-step recombination procedure (Skerker et al., 2005). |
| NABC724 | CB15N; *cada2* (R114A); $P_{ccna\_00746}$-*YFP::kan<sup>R</sup>* | NABC723 strain was transformed with pNABC735 plasmid to integrate $P_{ccna\_00746}$-YFP at the *xyl* locus. |
| NABC725 | CB15N; Δ*cada2*; pMT463-$P_{xyl}$-*cada2-flag::gent<sup>R</sup>* | NABC708 was transformed with pNABC747 plasmid. |
| NABC726 | CB15N; Δ*cada2*; pMT463-$P_{xyl}$-*cada2*(R68A)-*flag::gent<sup>R</sup>* | NABC708 was transformed with pNABC748 plasmid. |
| NABC727 | CB15N; Δ*cada2*; pMT687-$P_{xyl}$-*cada2*(R114A)-*flag::gent<sup>R</sup>* | NABC708 was transformed with pNABC749 plasmid. |
| NABC728 | CB15N; Δ*cada2*; pMT687-$P_{xyl}$-*cada2*(C267A)-*flag::gent<sup>R</sup>* | NABC708 was transformed with pNABC750 plasmid. |
| NABC729 | CB15N; Δ*cada2*; pMT687-$P_{xyl}$-*cada2<sup>myxococcus</sup>*::*gent<sup>R</sup>* | NABC708 was transformed with pMT463-pNABC751 plasmid. |
| NABC730 | CB15N; Δ*cada2*; pMT687-$P_{xyl}$-*cada2<sup>myxococcus</sup>*::*gent<sup>R</sup>*; $P_{ccna\_00746}$-*YFP::kan<sup>R</sup>* | NABC711 was transformed with pNABC751 plasmid. |
| NABC731 | CB15N; Δ*cada2*; pMT687-$P_{xyl}$-*EcAda*::*gent<sup>R</sup>*; $P_{ccna\_00746}$-*YFP::kan<sup>R</sup>* | NABC711 was transformed with pNABC752 plasmid. |
| NABC732 | CB15N; Δ*cada2*; $P_{EcAda}$-*YFP::kan<sup>R</sup>* | NABC708 strain was transformed with pNABC753 plasmid to integrate $P_{EcAda}$-YFP at the *xyl* locus. |
| NABC733 | CB15N; Δ*cada2*; pMT687-$P_{xyl}$-*cada2<sup>myxococcus</sup>*::*gent<sup>R</sup>*; $P_{EcAda}$-*YFP::kan<sup>R</sup>* | NABC732 was transformed with pNABC751 plasmid. |
| NABC734 | CB15N; Δ*cada2*; pMT687-$P_{xyl}$-*EcAda*::*gent<sup>R</sup>*; $P_{EcAda}$-*YFP::kan<sup>R</sup>* | NABC732 was transformed with pNABC752 plasmid. |
| NABC736 | BTH101; pKT25-rpoA::kan<sup>R</sup>; pUT18C-cada2::carb<sup>R</sup> | BTH101 was co-transformed with pNABC754 and pNABC755 for the bacterial-two-hybrid assay |
| NABC737 | BTH101; pKT25-rpoB::kan<sup>R</sup>; pUT18C-cada2::carb<sup>R</sup> | BTH101 was co-transformed with pNABC754 and pNABC756 for the bacterial-two-hybrid assay |
| NABC738 | BTH101; pKT25-rpoC::kan<sup>R</sup>; pUT18C-cada2::carb<sup>R</sup> | BTH101 was co-transformed with pNABC754 and pNABC757 for the bacterial-two-hybrid assay |
| NABC739 | BTH101; pKT25-rpoD::kan<sup>R</sup>; pUT18C-cada2::carb<sup>R</sup> | BTH101 was co-transformed with pNABC754 and pNABC758 for the bacterial-two-hybrid assay |
| NABC740 | BTH101; pKT25-rpoZ::kan<sup>R</sup>; pUT18C-cada2::carb<sup>R</sup> | BTH101 was co-transformed with pNABC754 and pNABC759 for the bacterial-two-hybrid assay |

**Table S2:**
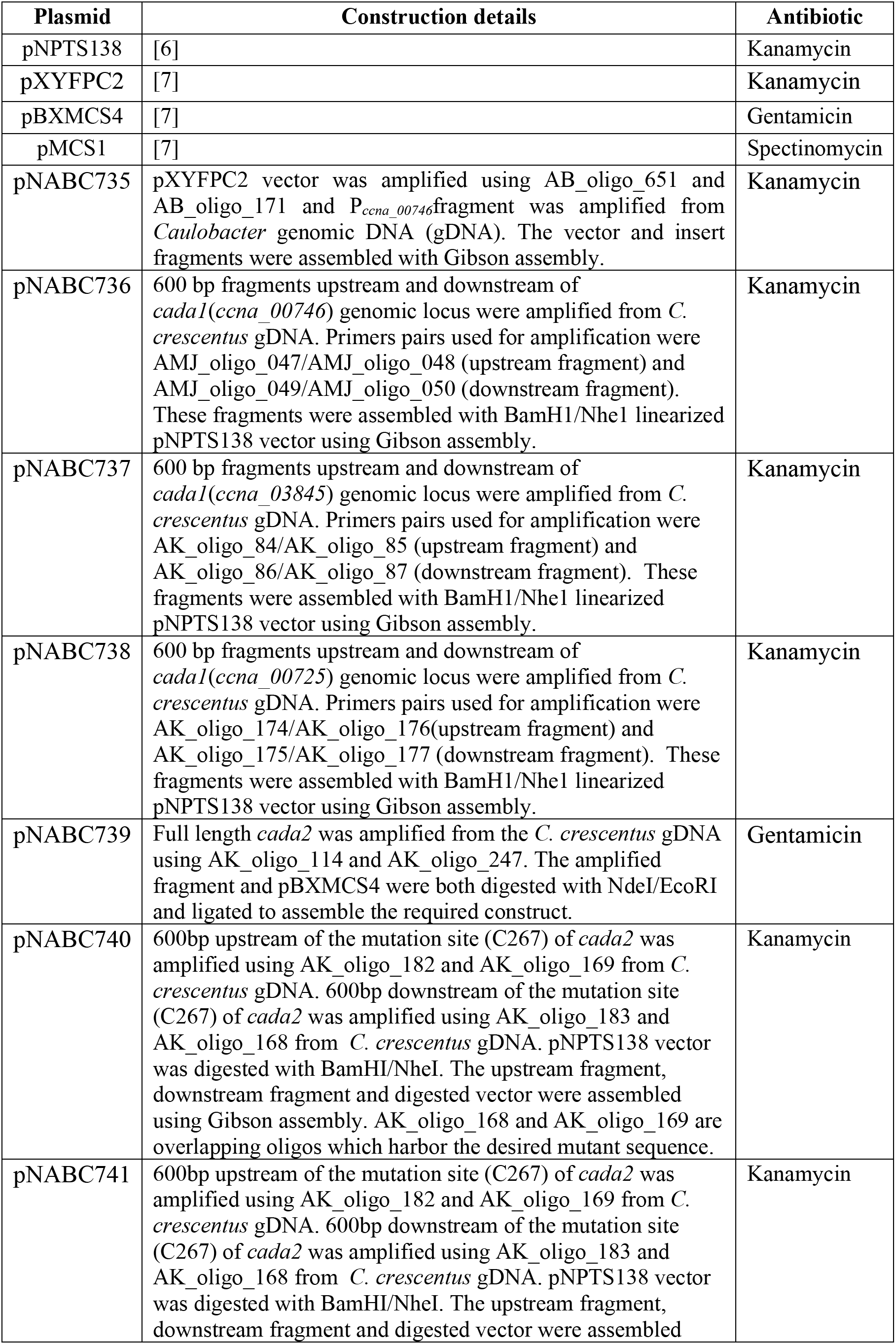
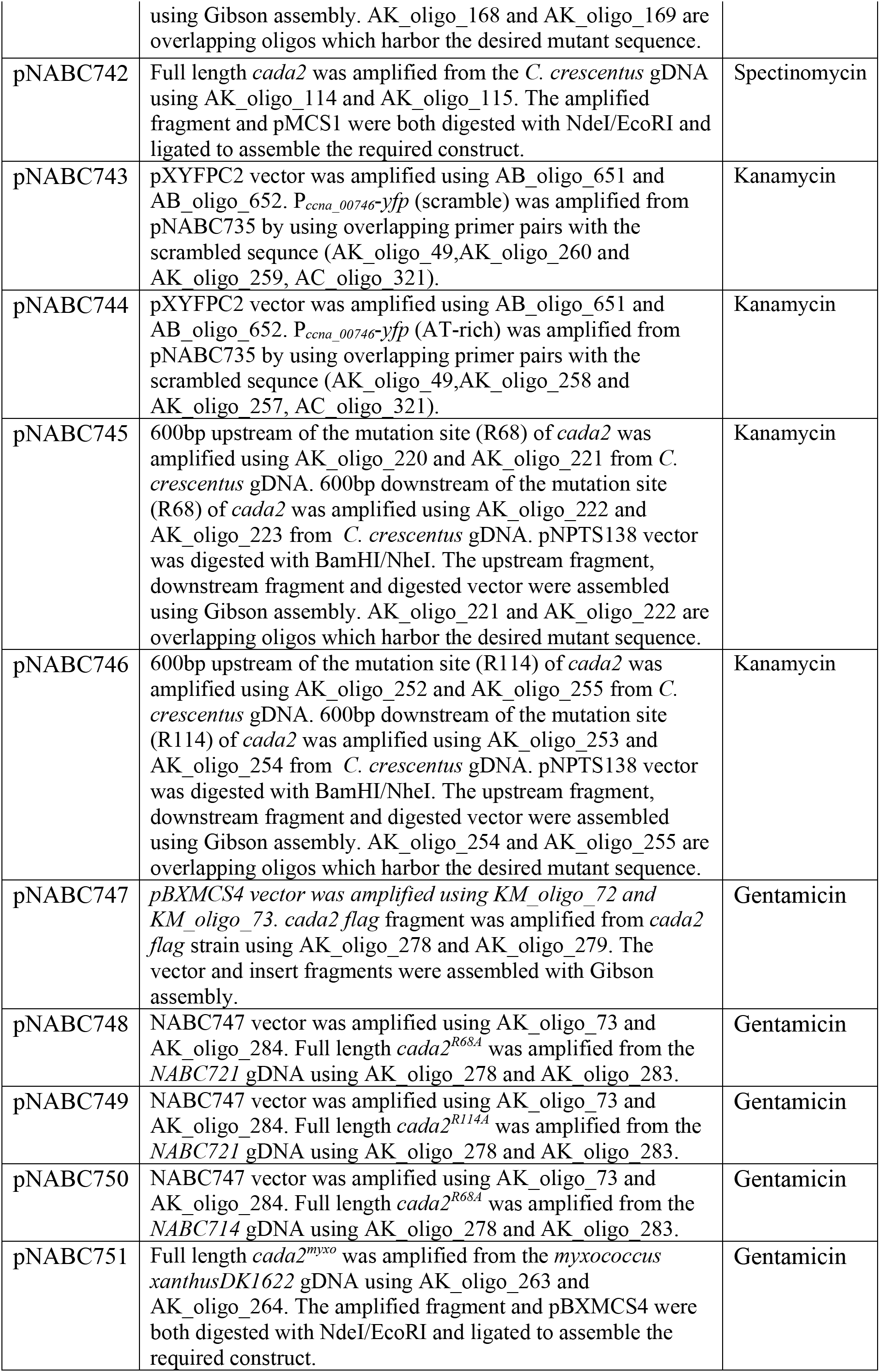
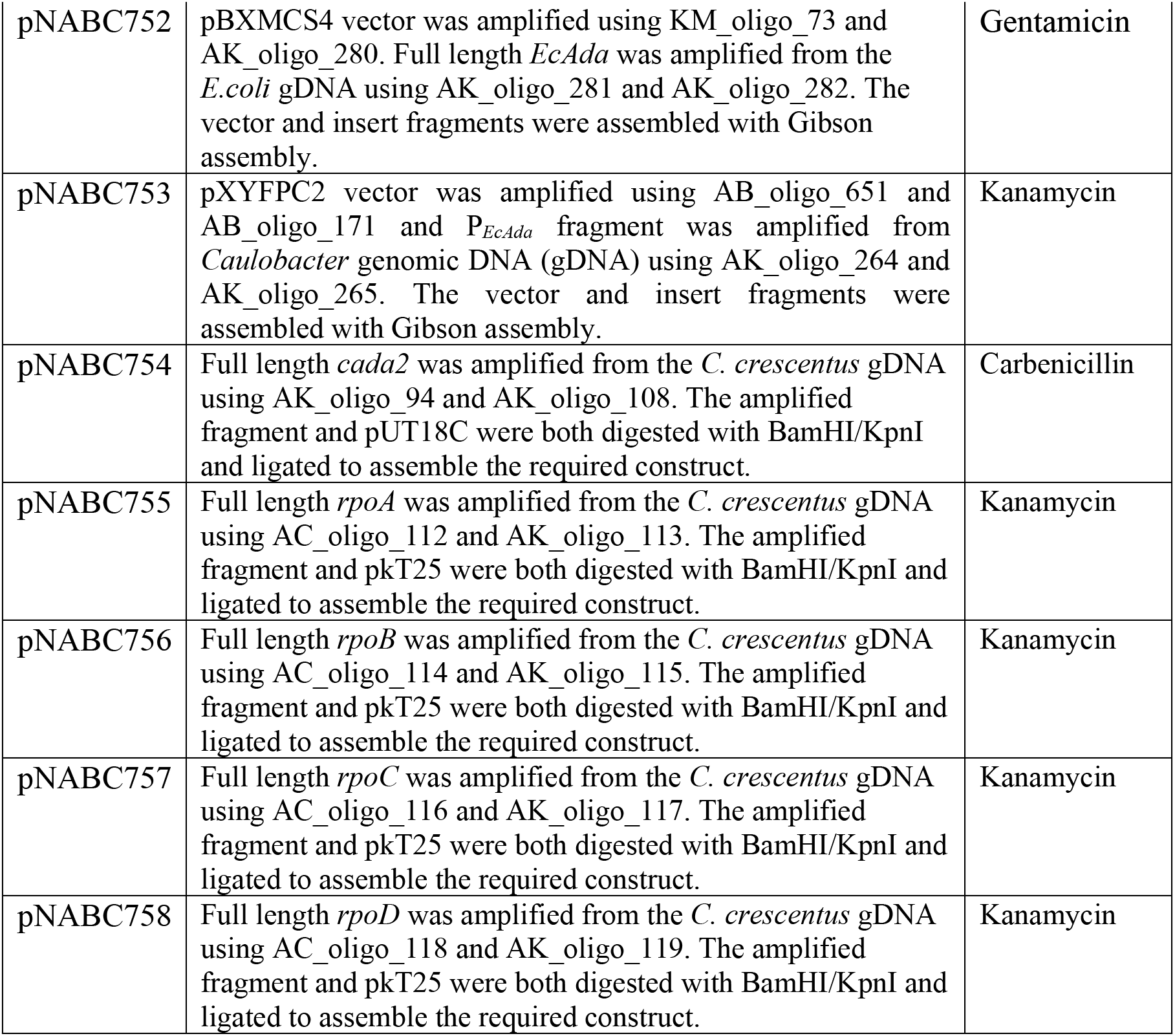
Plasmids used in present study.

| Plasmid | Construction details | Antibiotic |
| --- | --- | --- |
| pNPTS138 | [6] | Kanamycin |
| pXYFPC2 | [7] | Kanamycin |
| pBXMCS4 | [7] | Gentamicin |
| pMCS1 | [7] | Spectinomycin |
| pNABC735 | pXYFPC2 vector was amplified using AB_oligo_651 and AB_oligo_171 and P<sub>ccna_00746</sub> fragment was amplified from *Caulobacter* genomic DNA (gDNA). The vector and insert fragments were assembled with Gibson assembly. | Kanamycin |
| pNABC736 | 600 bp fragments upstream and downstream of *cada1(ccna_00746)* genomic locus were amplified from *C. crescentus* gDNA. Primers pairs used for amplification were AMJ_oligo_047/AMJ_oligo_048 (upstream fragment) and AMJ_oligo_049/AMJ_oligo_050 (downstream fragment). These fragments were assembled with BamHI/NheI linearized pNPTS138 vector using Gibson assembly. | Kanamycin |
| pNABC737 | 600 bp fragments upstream and downstream of *cada1(ccna_03845)* genomic locus were amplified from *C. crescentus* gDNA. Primers pairs used for amplification were AK_oligo_84/AK_oligo_85 (upstream fragment) and AK_oligo_86/AK_oligo_87 (downstream fragment). These fragments were assembled with BamHI/NheI linearized pNPTS138 vector using Gibson assembly. | Kanamycin |
| pNABC738 | 600 bp fragments upstream and downstream of *cada1(ccna_00725)* genomic locus were amplified from *C. crescentus* gDNA. Primers pairs used for amplification were AK_oligo_174/AK_oligo_176 (upstream fragment) and AK_oligo_175/AK_oligo_177 (downstream fragment). These fragments were assembled with BamHI/NheI linearized pNPTS138 vector using Gibson assembly. | Kanamycin |
| pNABC739 | Full length *cada2* was amplified from the *C. crescentus* gDNA using AK_oligo_114 and AK_oligo_247. The amplified fragment and pBXMCS4 were both digested with NdeI/EcoRI and ligated to assemble the required construct. | Gentamicin |
| pNABC740 | 600bp upstream of the mutation site (C267) of *cada2* was amplified using AK_oligo_182 and AK_oligo_169 from *C. crescentus* gDNA. 600bp downstream of the mutation site (C267) of *cada2* was amplified using AK_oligo_183 and AK_oligo_168 from *C. crescentus* gDNA. pNPTS138 vector was digested with BamHI/NheI. The upstream fragment, downstream fragment and digested vector were assembled using Gibson assembly. AK_oligo_168 and AK_oligo_169 are overlapping oligos which harbor the desired mutant sequence. | Kanamycin |
| pNABC741 | 600bp upstream of the mutation site (C267) of *cada2* was amplified using AK_oligo_182 and AK_oligo_169 from *C. crescentus* gDNA. 600bp downstream of the mutation site (C267) of *cada2* was amplified using AK_oligo_183 and AK_oligo_168 from *C. crescentus* gDNA. pNPTS138 vector was digested with BamHI/NheI. The upstream fragment, downstream fragment and digested vector were assembled | Kanamycin |
|  | using Gibson assembly. AK_oligo_168 and AK_oligo_169 are overlapping oligos which harbor the desired mutant sequence. |  |
| pNABC742 | Full length *cada2* was amplified from the *C. crescentus* gDNA using AK_oligo_114 and AK_oligo_115. The amplified fragment and pMCS1 were both digested with NdeI/EcoRI and ligated to assemble the required construct. | Spectinomycin |
| pNABC743 | pXYFPC2 vector was amplified using AB_oligo_651 and AB_oligo_652. *P<sub>ccna_00746</sub>-yfp* (scramble) was amplified from pNABC735 by using overlapping primer pairs with the scrambled sequence (AK_oligo_49, AK_oligo_260 and AK_oligo_259, AC_oligo_321). | Kanamycin |
| pNABC744 | pXYFPC2 vector was amplified using AB_oligo_651 and AB_oligo_652. *P<sub>ccna_00746</sub>-yfp* (AT-rich) was amplified from pNABC735 by using overlapping primer pairs with the scrambled sequence (AK_oligo_49, AK_oligo_258 and AK_oligo_257, AC_oligo_321). | Kanamycin |
| pNABC745 | 600bp upstream of the mutation site (R68) of *cada2* was amplified using AK_oligo_220 and AK_oligo_221 from *C. crescentus* gDNA. 600bp downstream of the mutation site (R68) of *cada2* was amplified using AK_oligo_222 and AK_oligo_223 from *C. crescentus* gDNA. pNPTS138 vector was digested with BamHI/NheI. The upstream fragment, downstream fragment and digested vector were assembled using Gibson assembly. AK_oligo_221 and AK_oligo_222 are overlapping oligos which harbor the desired mutant sequence. | Kanamycin |
| pNABC746 | 600bp upstream of the mutation site (R114) of *cada2* was amplified using AK_oligo_252 and AK_oligo_255 from *C. crescentus* gDNA. 600bp downstream of the mutation site (R114) of *cada2* was amplified using AK_oligo_253 and AK_oligo_254 from *C. crescentus* gDNA. pNPTS138 vector was digested with BamHI/NheI. The upstream fragment, downstream fragment and digested vector were assembled using Gibson assembly. AK_oligo_254 and AK_oligo_255 are overlapping oligos which harbor the desired mutant sequence. | Kanamycin |
| pNABC747 | *pBXMCS4* vector was amplified using KM_oligo_72 and KM_oligo_73. *cada2* flag fragment was amplified from *cada2* flag strain using AK_oligo_278 and AK_oligo_279. The vector and insert fragments were assembled with Gibson assembly. | Gentamicin |
| pNABC748 | NABC747 vector was amplified using AK_oligo_73 and AK_oligo_284. Full length *cada2*<sup>R68A</sup> was amplified from the NABC721 gDNA using AK_oligo_278 and AK_oligo_283. | Gentamicin |
| pNABC749 | NABC747 vector was amplified using AK_oligo_73 and AK_oligo_284. Full length *cada2*<sup>R114A</sup> was amplified from the NABC721 gDNA using AK_oligo_278 and AK_oligo_283. | Gentamicin |
| pNABC750 | NABC747 vector was amplified using AK_oligo_73 and AK_oligo_284. Full length *cada2*<sup>R68A</sup> was amplified from the NABC714 gDNA using AK_oligo_278 and AK_oligo_283. | Gentamicin |
| pNABC751 | Full length *cada2*<sup>myxo</sup> was amplified from the *myxococcus xanthus* DK1622 gDNA using AK_oligo_263 and AK_oligo_264. The amplified fragment and pBXMCS4 were both digested with NdeI/EcoRI and ligated to assemble the required construct. | Gentamicin |
| pNABC752 | pBXMCS4 vector was amplified using KM_oligo_73 and AK_oligo_280. Full length *EcAda* was amplified from the *E.coli* gDNA using AK_oligo_281 and AK_oligo_282. The vector and insert fragments were assembled with Gibson assembly. | Gentamicin |
| pNABC753 | pXYFPC2 vector was amplified using AB_oligo_651 and AB_oligo_171 and P<sub>EcAda</sub> fragment was amplified from *Caulobacter* genomic DNA (gDNA) using AK_oligo_264 and AK_oligo_265. The vector and insert fragments were assembled with Gibson assembly. | Kanamycin |
| pNABC754 | Full length *cada2* was amplified from the *C. crescentus* gDNA using AK_oligo_94 and AK_oligo_108. The amplified fragment and pUT18C were both digested with BamHI/KpnI and ligated to assemble the required construct. | Carbenicillin |
| pNABC755 | Full length *rpoA* was amplified from the *C. crescentus* gDNA using AC_oligo_112 and AK_oligo_113. The amplified fragment and pkT25 were both digested with BamHI/KpnI and ligated to assemble the required construct. | Kanamycin |
| pNABC756 | Full length *rpoB* was amplified from the *C. crescentus* gDNA using AC_oligo_114 and AK_oligo_115. The amplified fragment and pkT25 were both digested with BamHI/KpnI and ligated to assemble the required construct. | Kanamycin |
| pNABC757 | Full length *rpoC* was amplified from the *C. crescentus* gDNA using AC_oligo_116 and AK_oligo_117. The amplified fragment and pkT25 were both digested with BamHI/KpnI and ligated to assemble the required construct. | Kanamycin |
| pNABC758 | Full length *rpoD* was amplified from the *C. crescentus* gDNA using AC_oligo_118 and AK_oligo_119. The amplified fragment and pkT25 were both digested with BamHI/KpnI and ligated to assemble the required construct. | Kanamycin |

**Table S3:**
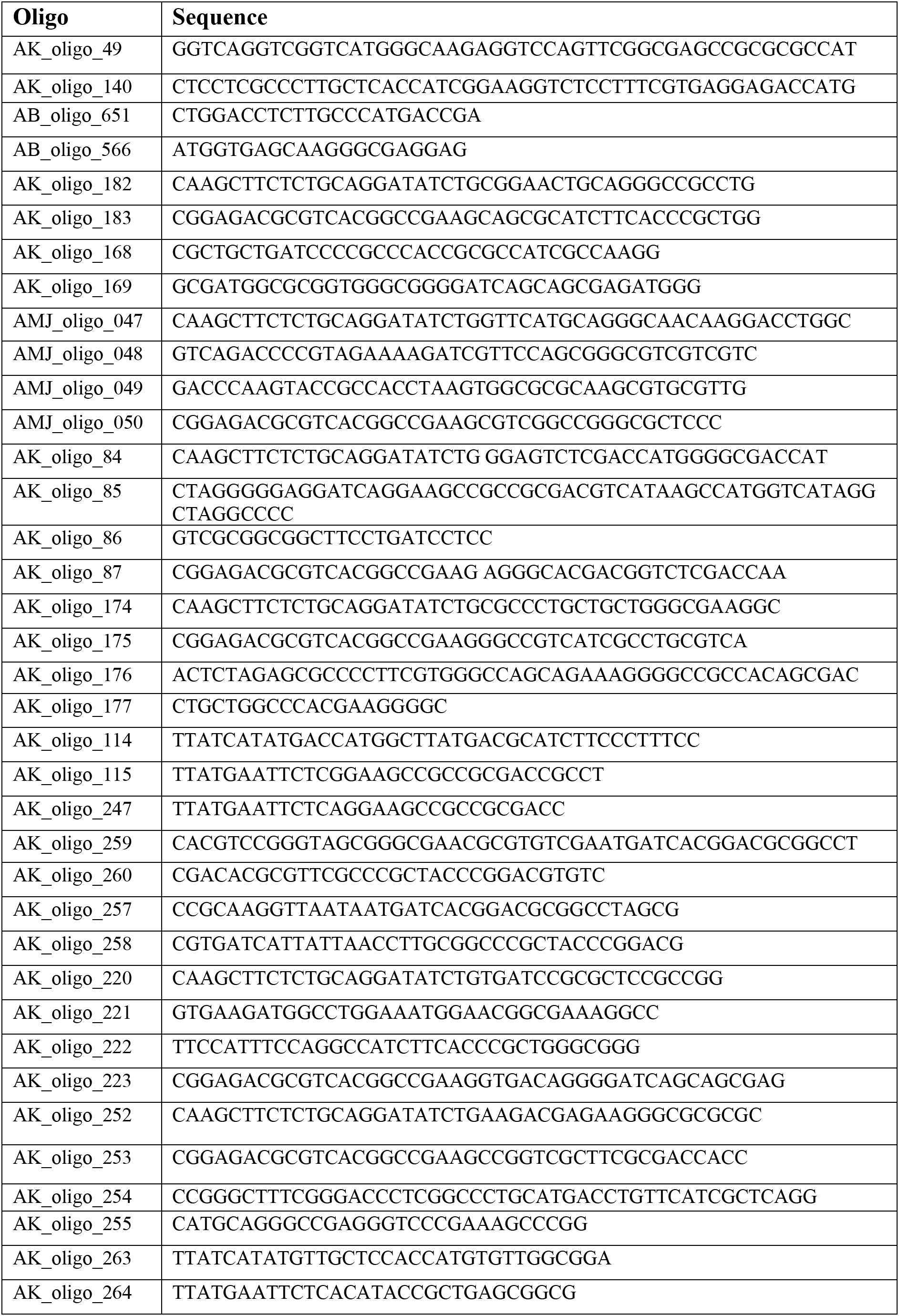
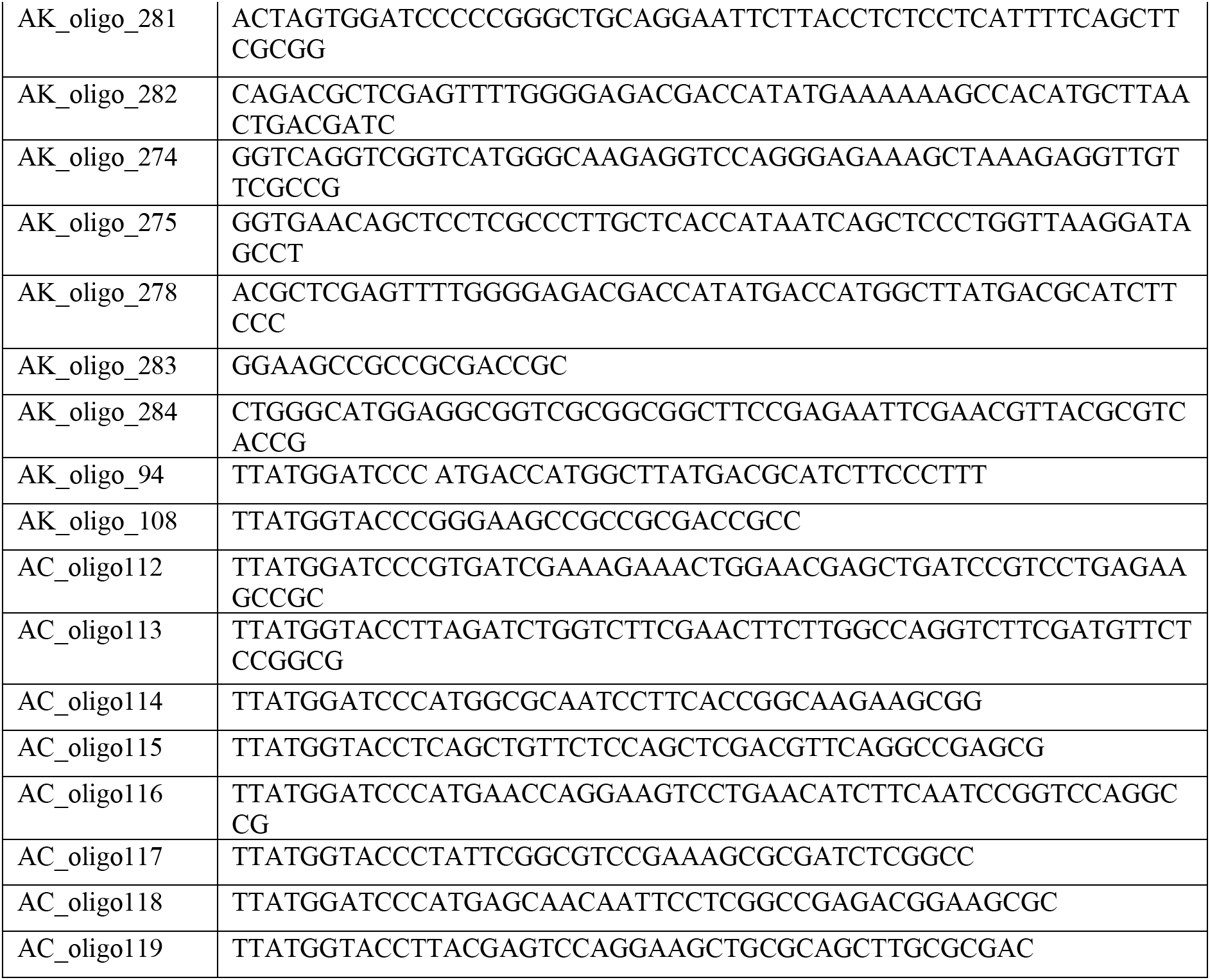
Oligos used in present study.

